# *Cis*-membrane association of human ATG8 proteins N-terminus mediates autophagy

**DOI:** 10.1101/2022.06.10.495627

**Authors:** Wenxin Zhang, Taki Nishimura, Deepanshi Gahlot, Chieko Saito, Colin Davis, Harold B. J. Jefferies, Anne Schreiber, Lipi Thukral, Sharon A. Tooze

## Abstract

Autophagy is an essential catabolic pathway which sequesters and engulfs cytosolic substrates via autophagosomes, unique double-membraned structures. ATG8 proteins are ubiquitin-like proteins recruited to autophagosome membranes by lipidation at the C-terminus. ATG8s recruit substrates, such as p62, and play an important role in mediating autophagosome membrane expansion. However, the precise function of lipidated ATG8 in expansion remains obscure. Using a real-time *in vitro* lipidation assay, we revealed that the N-termini of lipidated human ATG8s (LC3B and GABARAP) are highly dynamic and interact with the membrane. Moreover, atomistic MD simulation and FRET assays indicate that N-termini of LC3B and GABARAP associate *in cis* on the membrane. The *cis*-membrane association of the N-terminus is critical to maintain membrane expansion and the size of autophagosomes in cells, consequently, mediating the efficient degradation of p62. Our study provides fundamental molecular insights into autophagosome membrane expansion, revealing the critical and unique function of lipidated ATG8.

## Introduction

Macroautophagy (hereafter autophagy) is a bulk intracellular degradation pathway, in which long-lived or damaged cytoplasmic materials are enveloped by double membraned organelles, called autophagosomes, and delivered to lysosomes. Autophagy occurs in both basal and stressed conditions to maintain cellular homeostasis and cell survival (Green and Levine, 2014; Levine and Kroemer, 2008). Upon autophagy induction, a small membrane structure, a phagophore, grows and expands becoming a unique cup-shaped structure. Subsequently, the open edges of the phagophore close to form a spherical mature autophagosome (Nakatogawa, 2020). Autophagy is a highly dynamic membrane process and sources lipids from multiple cellular membrane compartments, including ER, mitochondria, MAM (mitochondria associated ER membrane), Golgi, ERES (ER exit sites), ERGIC (ER-Golgi intermediate compartment) and the plasma membrane (Nishimura and Tooze, 2020; Tooze and Yoshimori, 2010).

ATG8 is a unique ubiquitin-like protein conjugated to phospholipids on autophagic membranes (Ichimura et al., 2000; Kabeya et al., 2004). In yeast, there is only one Atg8, while in mammals there are at least six distinct ATG8 proteins, classified into two subfamilies, LC3s (LC3A, LC3B and LC3C) and GABARAPs (GABARAP, GABARAPL1 and GABARAPL2) (Mizushima, 2020). These proteins are widely used as markers of autophagosomes. The conjugation reaction starts with the cleavage of ATG8 by ATG4 to expose a glycine residue at the C-terminus. Subsequently, ATG8 is activated by ATG7 (E1) and transferred to ATG3 (E2), and finally covalently conjugated to phosphatidylethanolamine (PE) with the assistance of the ATG12–ATG5-ATG16L1 complex (hereafter, the E3 complex) (Martens and Fracchiolla, 2020). These lipidated ATG8 proteins on autophagic membranes function as adaptors to recruit cytosolic receptors, such as p62/SQSTM1, NBR1 and optineurin. These ATG8 interactors contain a consensus motif, called the Atg8-interacting motif (AIM) or the LC3-interacting region (LIR), which docks into two hydrophobic pockets on the surface of ATG8s (Johansen and Lamark, 2020; Stolz et al., 2014).

In addition to selectively binding cytosolic cargo, lipidated ATG8 proteins play an important role in mediating phagophore (also called isolation membranes) growth (Xie et al., 2008). *In vivo* evidence in cells suggests that ATG8 family proteins contribute to phagophore membrane expansion. Reduction in the protein level of Atg8 decreases the size of the autophagic bodies in yeast (Xie et al., 2008) and autophagosome size during *C. elegans* embryogenesis (Wu et al., 2015). In line with this, in mammalian cells, knock out of all ATG8 proteins results in smaller autophagosomes, though autophagosome formation is still maintained (Nguyen et al., 2016). Several *in vitro* studies have focused on the role of lipidated ATG8s in membrane tethering and fusion. The first studies employed *in vitro* reconstitution with yeast Atg7, Atg3 and Atg8 to study Atg8-PE and suggested Atg8-PE contributes to phagophore expansion by mediating hemifusion (Nakatogawa et al., 2007). Similar experiments have been employed to study mammalian ATG8 proteins, using either ATG8s chemically conjugated to membranes or lipidated ATG8s to demonstrate the fusogenic properties of ATG8s (Landajuela et al., 2016; Taniguchi et al., 2020; Weidberg et al., 2011). Although these studies show association between lipidated ATG8 proteins on two distinct lipid bilayers (“*in trans*”), a recent study demonstrated that two-aromatic membrane facing residues of Atg8 can associate with the membrane on which Atg8 is lipidated (“*in cis”*) (Maruyama et al., 2021). Yet, the underlying mechanism of ATG8-dependent phagophore expansion and the function of ATG8s in autophagy remain to be fully understood.

Crucially, unlike ubiquitin and other ubiquitin-like proteins, ATG8 family proteins contain additional two a-helices at their N-termini, which is an evolutionary conserved feature (Wesch et al., 2020). Previous studies have reported several distinct functions of the N-terminal regions, such as p62 recognition (Shvets et al., 2011; Shvets et al., 2008), membrane tethering (Nakatogawa et al., 2007; Weidberg et al., 2011; Wu et al., 2015), oligomerisation (Coyle et al., 2002; Nakatogawa et al., 2007) and direct lipid binding (Chu et al., 2013; Sentelle et al., 2012). These versatile roles might be achieved by the structural flexibility of ATG8 N-terminus (Wu et al., 2015).

Here we determine the conformation dynamics of LC3B and GABARAP N-terminal regions during lipidation using a number of approaches including a real time assay. We show that the N-termini of LC3B and GABARAP not only contribute to the interaction with ATG7 and ATG3 during the conjugation reaction, but also associate with the membrane after their lipidation. Interestingly, these regions associate in *cis* with the membrane where LC3B and GABARAP are lipidated. *In vitro* FRET assays and molecular dynamics (MD) simulation analyses confirmed the *cis* membrane interaction of LC3B/GABARAP N-termini. Furthermore, cells expressing GABARAP mutants, lacking N-terminal regions or with impaired *cis*-membrane interactions of the N-terminus, have small autophagosome-like structures and abnormal p62 degradation. Our data support a model that efficient autophagic membrane expansion and p62 degradation requires the N-termini of the ATG8 proteins. The association of the N-terminus with membranes *in cis* coordinates lipidation and membrane expansion, and provides a molecular mechanism underlying the unique requirement amongst all the ubiquitin-like proteins for ATG8s in autophagy.

## Results

### N-terminal regions of LC3B/GABARAP are dynamically incorporated into hydrophobic environments during in vitro lipidation reaction

To investigate how the N-terminal regions of LC3B/GABARAP behave during the conjugation and lipidation reaction, we developed a real-time *in vitro* assay using LC3B/GABARAP proteins labelling with 7-nitrobenz-2-oxa-1,3-diazol-4-yl (NBD) (**Figure 1A**). As NBD is an environmentally sensitive fluorescence probe that displays enhanced fluorescence around 535 nm in a hydrophobic environment, thus increases in NBD fluorescence reflect the location of LC3B/GABARAP N-termini in protein-protein and protein-membrane interfaces. LC3B/GABARAP were labelled with NBD by introducing single mutations at N-terminal residues: LC3B S3C^NBD^ and GABARAP K2C^NBD^, respectively. We first mixed ATG7, ATG3, the E3 complex, LC3B S3C^NBD^ or GABARAP K2C^NBD^ and liposomes. After the addition of ATP, the increase in NBD signal was measured by fluorescence spectroscopy. As shown in **Figure 1B**, the fluorescence of LC3B S3C^NBD^ was significantly increased in the presence of all components essential for efficient LC3 lipidation reaction (“All”). In contrast, no increase in fluorescence was observed without the E1 enzyme ATG7 (“No ATG7”), suggesting that the increase in NBD fluorescence is caused by the LC3 lipidation reaction. We observed a partial increase of the NBD fluorescence even in the absence of the E2 enzyme ATG3 (“No ATG3”) or liposomes (“No liposomes”) (**Figure 1B**), in which ATG7∼LC3B and/or ATG3∼LC3B intermediates, but not LC3B-PE, are formed. Furthermore, ATG7 and ATP were sufficient to induce a partial increase of NBD signals, which was enhanced by the addition of ATG3 (**Figure 1-figure supplement 1A and 1B**). Therefore, the NBD fluorescence increase reflects the formation of ATG7∼LC3B and ATG3∼LC3B intermediates as well as that of LC3B-PE (**Figure 1A**). We biochemically confirmed that LC3B S3C^NBD^ was properly conjugated to ATG7, ATG3 and PE in *in vitro* reactions (**Figure 1C**). Similar results were acquired using GABARAP K2C^NBD^ (**Figure 1D and 1E; Figure 1-figure supplement 1A and 1B**). These results suggest that the N-terminal regions of LC3B/GABARAP are incorporated into a hydrophobic environment during the conjugation and lipidation reaction.

**Figure 1.**
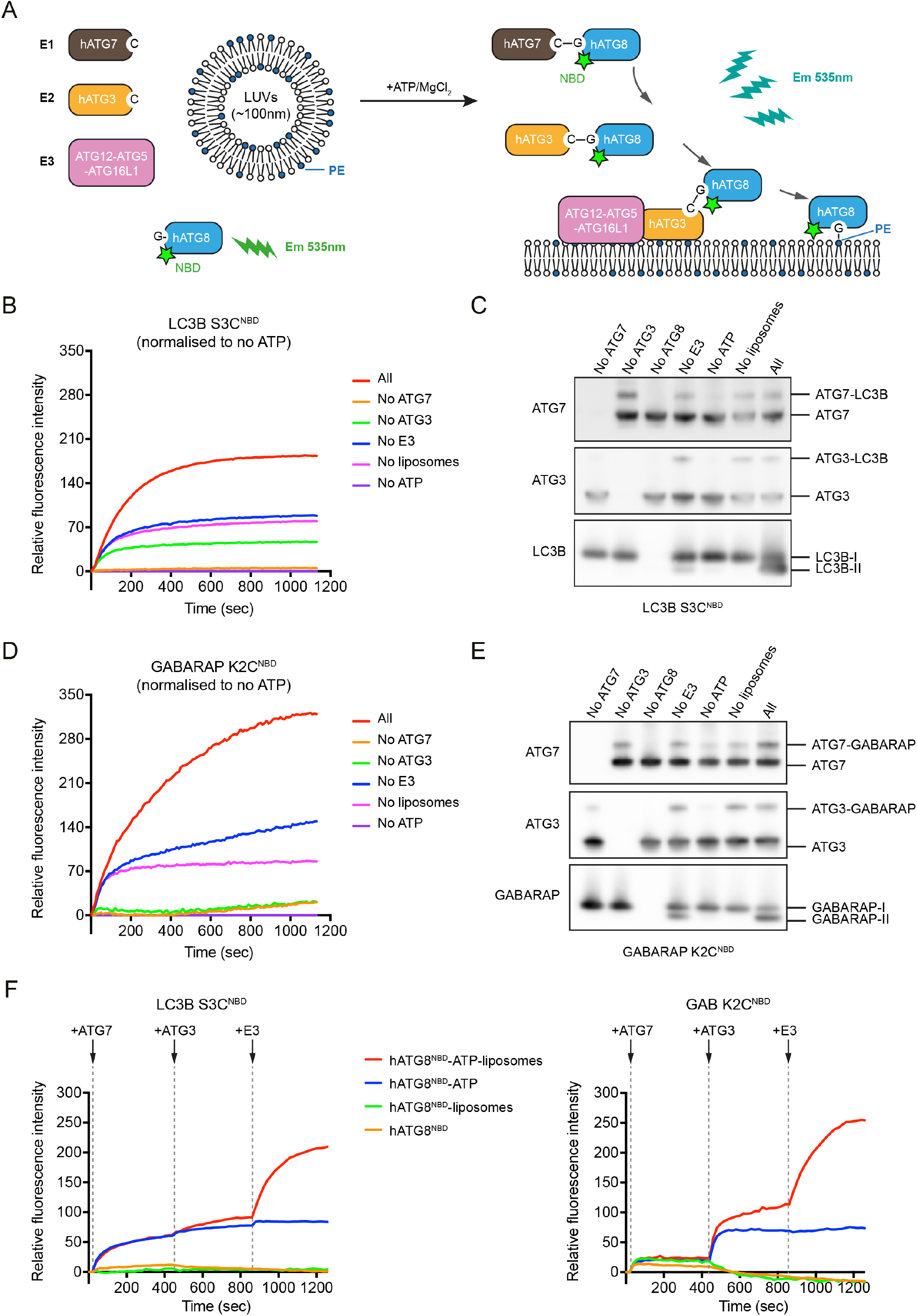
A real-time assay to track LC3B and GABARAP N-termini dynamics during lipidation. **(A)** A schematic diagram of the real time lipidation assay using N-terminal NBD labelled hATG8. **(B)** The NBD fluorescence changes of LC3B S3C^NBD^ during lipidation reaction. “All” condition contains 0.2 μM ATG7, 0.2 μM ATG3, 0.05 μM ATG12–ATG5-ATG16L1, 1 mM ATP, 1 mM LUVs (50% DOPE /50% POPC) and 1 μM LC3B S3C^NBD^, at 37°C. The NBD fluorescence was traced immediately once ATP was added (time point at 0s). The rest conditions were performed in the absence of one component from the “All” condition. The relative NBD fluorescence was normalised to “no ATP” condition. Data represent mean values (n=3). **(C)** Western blots of samples from **(B)**. After 20 min reaction, the reactions from the real-time assay were stopped by adding sample buffer and immunoblot for ATG7, ATG3 and LC3B. **(D**) The NBD fluorescence changes of GABARAP K2C^NBD^ during lipidation reaction. Experimental conditions are the same as described in **(B)**. Data represent mean values (n=3). **(E)** Western blots of samples from **(D). (F)** Step by step real-time lipidation assay with LC3B S3C^NBD^ and GABARAP K2C^NBD^. The NBD fluorescence was recorded continuously while adding ATG7 (between 60-80s), adding ATG3 (between 480-500s) and adding E3 complex (between 900-920s). Data represent mean values (n=3).

To dissect the effects of intermediate formation and lipidation on the increase in NBD fluorescence, we employed the real-time assay and added each component sequentially. The NBD signal of LC3B S3C^NBD^ increased within 30 sec after the addition of ATG7, independently of liposomes. After that, the E3 complex strongly stimulated the NBD signal increase in the presence of liposomes (**Figure 1F**, left). In the case of GABARAP K2C^NBD^, similar responses of NBD signals were observed after the addition of ATG7, ATG3 and the E3 complex, while ATG3 more efficiently increased the NBD signal compared to ATG7 (**Figure 1F**, right). Thus, these results indicate that the N-termini of LC3B and GABARAP are highly dynamic during the lipidation reaction and encounter weak and strong hydrophobic environments during intermediate formation and lipid-conjugation, respectively.

### Lipidated LC3B/GABARAP associate with membranes *in cis* analysed in silico

The real-time NBD assay demonstrated that the LC3B/GABARAP N-termini are located in hydrophobic surroundings after the lipidation reaction (**Figure 1F**), implying that the N-termini of lipidated LC3B/GABARAP might be inserted into a lipid bilayer. So far, the structures of non-lipidated ATG8 protein have been extensively studied by protein crystallisation techniques and NMR (Sora et al., 2020). These high-resolution structures have been employed in coarse-gained (CG) simulations which suggest that lipidated LC3B associates with membranes via a patch of basic residues (R68, R69, R70), β3 and the hydrophobic C-terminal region after β4 (Fas et al., 2021; Thukral et al., 2015) (**Figure 1-figure supplement 1C**). On the other hand, there are two studies resolving the structure of lipidated GABARAP (Ma et al., 2010) and lipidated yeast Atg8 (Maruyama et al., 2021) on lipid bilayer nanodiscs using NMR. The NMR structures suggest similar membrane-association regions in GABARAP and yeast Atg8, which are also demonstrated in the CG simulation studies. However, there are discrepancies between these studies which suggest different orientations of ATG8-PE on the membrane.

To further investigate the conformational rearrangements and structural dynamics of lipidated LC3B and GABARAP, we utilised atomistic MD simulations instead of static methods like molecular docking. Our structural analysis of membrane-embedded LC3B and GABARAP is based on six atomistic, explicit-water MD simulation trajectories with a length of 1 μs each. The initial starting structures for these simulations were prepared after extensive orientation variability calculations on membrane to obtain unbiased position analysis (**Figure 2A; see Methods**). We found that the orientation variability calculations and membrane-facing residues converge to a unique and common conformation. Based on three replicas, the probability of each residue-lipid distance contact was computed and their occupancy i.e., presence during the simulation was calculated (**Figure 2-figure supplement 1B and 1D)**.

**Figure 2.**
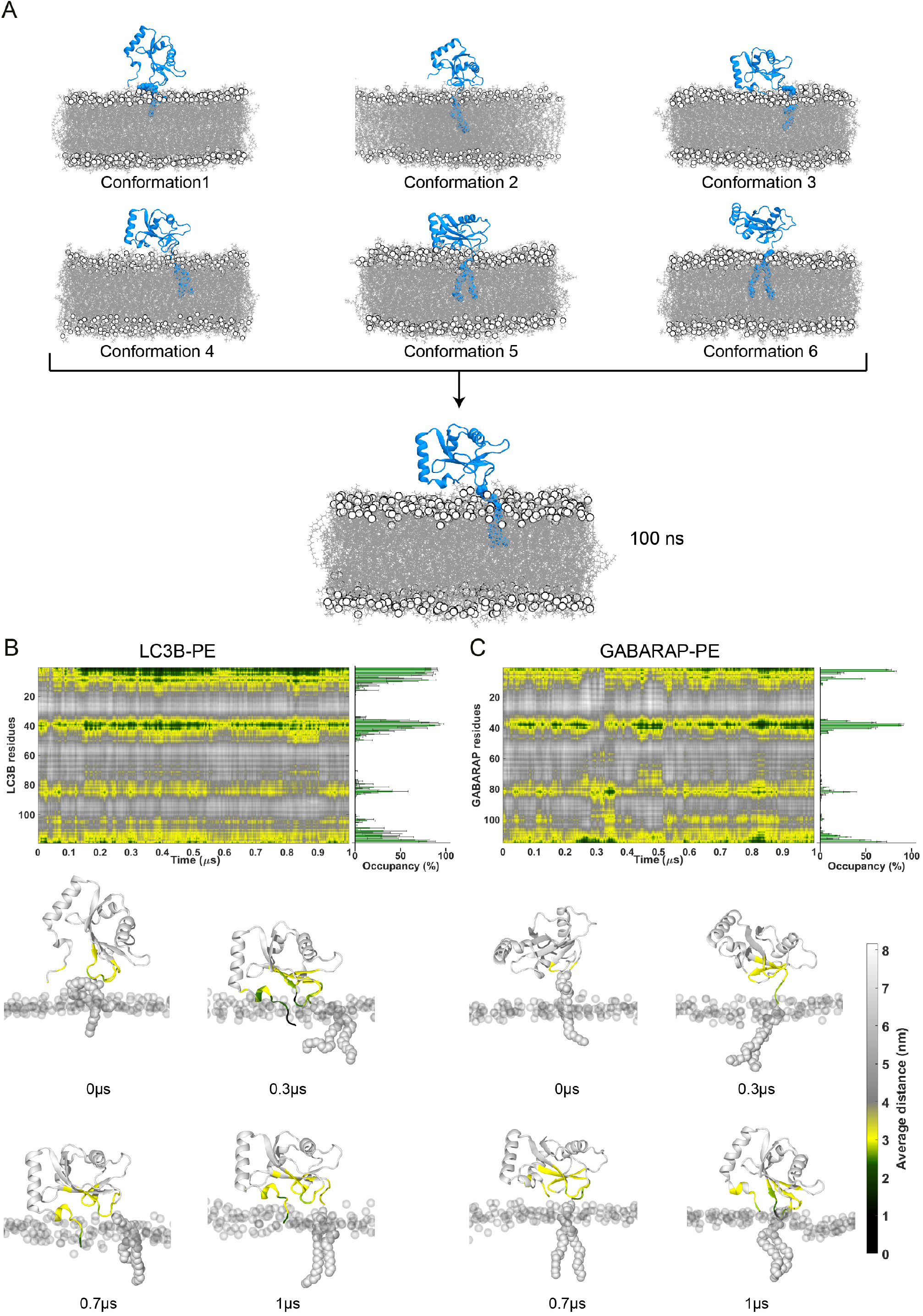
Molecular dynamics simulations of lipidated LC3B and GABARAP in POPC membrane. **(A)** Pool of six Initial conformations with varying lipid-contacting orientations. After an initial refinement of 100 ns, all conformations converged to a single orientation with N-terminal moving closer to lipids, **(B) and (C)** Time evolution of distance between LC3B/GABARAP residues respectively and POPC membrane highlighting four distinct membrane interacting regions mainly, N-terminal, loop 3, loop 6 and C-terminal. Horizontal bar graph in the right shows the contact occupancy (in percentage) of all replicates. The contacts were categorized into four parts depending upon the residues’ side chain orientation on the membrane: Inserted at ≤ 2 nm colored in black, at membrane surface at ≤ 2.8nm colored in green, in proximity at < 3 nm colored in yellow, and not interacting at > 3.5nm colored in grey. Lower panel depicts the snapshots at different time points, wherein protein residues are colored according to their distance with the membrane. The headgroup of POPC membrane is highlighted in transparent silver colour, whereas the protein and lipid anchor are highlighted in silver colour.

In lipidated LC3B and GABARAP, simulations revealed four prominent membrane-interacting interfaces: the N-termini, loop 3, loop 6 and C-termini were consistently interacting with membrane albeit with dynamic movements of the protein chain (**Figure 2B and 2C; Figure 2-figure supplement 1B and 1D**). We observed that the N-termini of LC3B/GABARAP were inserted into the membrane over the length of the simulations. Compared to lipidated GABARAP, there are more membrane-associated residues in lipidated LC3B (**Figure 2-figure supplement 1B and 1D**). Unlike previous CG-MD simulations (Fas et al., 2021; Thukral et al., 2015), the basic residues in LC3B (R68, R69, R70, hereafter RRR) or GABARAP (R65, K66, R68, hereafter RKR) were distant from the membrane and facing towards cytosol in our atomistic models. Recently, the aromatic residues F77/F79 in lipidated yeast Atg8 were shown to be inserted into the membrane and regulate the membrane deformation *in vitro* (Maruyama et al., 2021). However, our atomistic MD simulation results show that the corresponding residues F80 in LC3B and F77/F79 GABARAP do not associate with the membrane, while additional membrane contacts are found in loop 6. Interestingly, our MD simulation suggests a *cis* membrane-association model of lipidated LC3B/GABARAP. The simulations (**Figure 2B and 2C)** predict the N-termini of LC3B/GABARAP interacts with the same membrane where the ATG8-PE is inserted (**Figure 3A and 3B)**.

**Figure 3.**
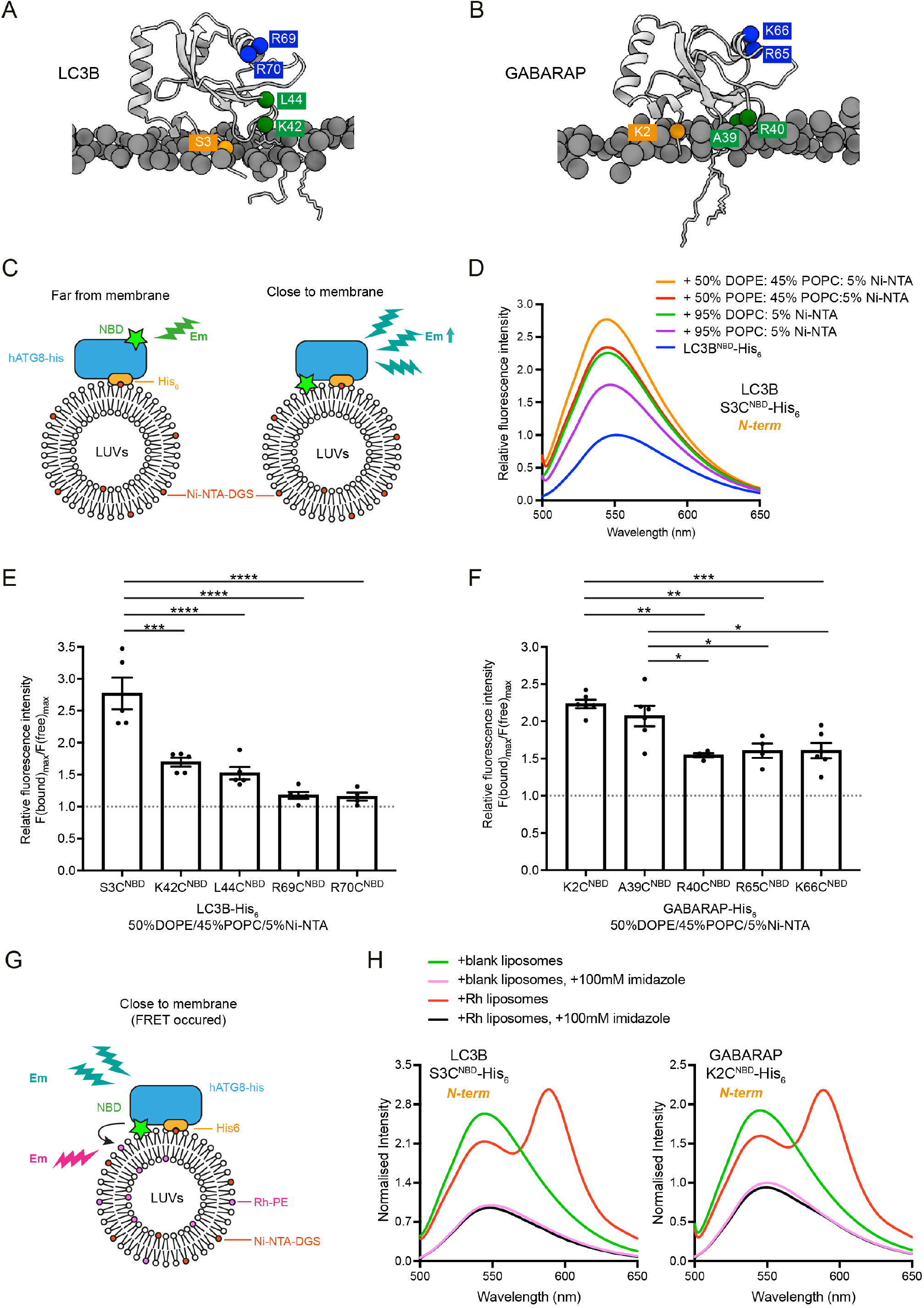
Analysis of membrane-association interface of liposome-conjugated LC3B and GABARAP. **(A-B)** Representative structures of lipidated LC3B (300 ns) and GABARAP (840 ns) from MD simulation. The individual amino acids of interest were labelled with NBD. **(C)** Scheme for the NBD spectra assay to characterise the membrane-association residues in LC3B and GABARAP, using LC3B-His_6_/GABARAP-His_6_ and LUVs containing nickel lipids to mimic the lipidated status. If the residue is embedded in the membrane, the NBD fluorescence gets increased. **(D)** Example of fluorescence spectra of LC3B S3C^NBD^-His_6_ in the absence of LUVs (protein only, 1 μM) and in the presence of LUVs containing 5% Ni-NTA (1 mM). Spectra represent mean values (n=5) **(E-F)** Quantification of NBD fluorescence increase of the screened residues in LC3B-His_6_ and GABARAP-His_6_. F(bound)_max_/F(free)_max_ ratio represents the maximum emission intensity of NBD-labelled LC3B-His_6_/GABARAP-His_6_ bound to liposomes containing 50% DOPE/45%POPC/5% Ni-NTA, normalised to the maximum emission intensity of NBD-labelled LC3B-His_6_/GABARAP-His_6_ in the absence of liposomes (n=4-5, mean ± SEM). (**G**) Schematic diagram for the FRET assay to confirm the interaction between NBD-labelled LC3B-His_6_/GABARAP-His_6_ and rhodamine labelled liposomes. If the residue interacts with membrane, the emission of NBD would excite the rhodamine on the liposomes. (**H**) FRET assay with S3C^NBD^-His_6_ and GABARAP K2C^NBD^-His_6_. Each NBD-labelled LC3B-His_6_/GABARAP-His_6_ (1μM) was mixed with 1mM blank liposomes (50% DOPE/45% POPC/5% Ni-NTA) (green) or rhodamine liposomes (50% DOPE/43%POPC/5% Ni-NTA/2% Rh-PE) (red). In parallel, addition of 100 mM imidazole to remove NBD-labelled LC3B-His_6_/GABARAP-His_6_ from liposomes was performed as negative controls. Spectra represent mean values (n=3). Differences were statistically analysed by one-way ANOVA and Turkey multiple comparison test. *p<0.05, **p<0.01, ***p<0.001, ****p<0.0001.

### N-terminal regions of liposome-conjugated LC3B/GABARAP are inserted into a lipid bilayer

To validate our model for membrane-association of lipidated LC3B, we examined if the N-terminus of membrane-conjugated LC3B is able to associate *in cis*. In brief, C-terminal His_6_-tagged LC3B S3C^NBD^ was mixed with liposomes containing nickel lipids and then the increases of NBD signals were analysed (**Figure 3C**). If the N-terminus of LC3B is close to membranes, thereby encountering a hydrophobic environment, the NBD signals are increased. When LC3B S3C^NBD^-His_6_ was mixed with POPC (1-palmitoyl-2-oleoyl-sn-glycero-3-phosphocholine)-based liposomes containing 5% Ni-NTA lipid, the fluorescence intensity of NBD was significantly increased compared to LC3B S3C^NBD^-His_6_ alone (**Figure 3D**). In addition, to investigate the effects of lipid saturation and PE on membrane-association of LC3B N-terminus, we tested another three lipid compositions. These liposomes all contain 5% Ni-NTA-lipid, supplemented with 95% DOPC (1,2-dioleoyl-*sn*-glycero-3-phosphocholine), 50% POPE (1-palmitoyl-2-oleoyl-sn-glycero-3-phosphoethanolamine)/ 45% POPC or 50% DOPE (1,2-dioleoyl-*sn*-glycero-3-phosphoethanolamine)/45% POPC, respectively. As shown in **Figure 3D**, NBD signals increased in the presence of unsaturated lipids and/or PE, indicating that lipid packing defects can facilitate membrane association of the LC3B N-terminus. We further investigated whether other regions of LC3B are involved in membrane association. K42 and L44 residues in loop 3 and R69 and R70 residues in RRR region were labelled with NBD. The fluorescence intensity of the NBD label from LC3B K42C^NBD^-His_6_ and LC3B L44C^NBD^-His_6_ was increased in the presence of liposomes containing nickel lipids, though the increase was less than half of the increase observed with LC3B S3C^NBD^-His_6_ (**Figure 3E**). In contrast, no fluorescence changes were detected with LC3B R69C^NBD^-His_6_ and LC3B R70C^NBD^-His_6_ (**Figure 3E; Figure 3-figure supplement 1A**). Consistent with the atomistic MD simulation, these results suggest that the N-terminal and loop 3 regions of lipidated LC3 are close to membranes compared to the RRR region. We performed the same assays with C-terminal His_6_-tagged and NBD-labelled GABARAPs to examine the membrane association of GABARAP bound to liposomes. Consistent with our atomistic MD, the N-terminus (K2C^NBD^) and loop 3 (A39C^NBD^, R40C^NBD^) of GABARAP associated with membranes (**Figure 3F; Figure 3-figure supplement 1B**). Surprisingly, in contrast to the MD simulation model, the RKR region (R65C^NBD^, K66C^NBD^) of GABARAP also showed NBD fluorescence increase.

To further validate that the increase of NBD fluorescence intensity results from protein-membrane association, we designed a FRET assay using NBD labelled LC3B-His_6_ or GABARAP-His_6_ proteins and Rhodamine-labelled liposomes (**Figure 3G**). If the selected residues **(Figure 3A and 3B)** interact with liposomes containing rhodamine lipids, energy transfer should be detected, where the excitation of NBD results in an increase in rhodamine signal emission at 583nm and a concomitant reduction of NBD signal at 535nm (Hui et al., 2011). As shown in **Figure 3H**, we detected FRET between LC3B S3C^NBD^-His_6_ or GABARAP K2C^NBD^-His_6_ and rhodamine liposomes (“+Rh liposomes”), compared to experiments with liposomes containing no rhodamine lipids (“+blank liposomes”). The increase in the NBD signal or FRET did not occur when LC3B S3C^NBD^-His_6_ or GABARAP K2C^NBD^-His_6_ was released with imidazole. FRET experiments were also performed with the other NBD labelled LC3B-His_6_ and GABARAP-His_6_ proteins (**Figure 3-figure supplement 2**).

Our FRET experiments are consistent with the fluorescence-based liposome binding assays. The *in vitro* experimental results are in line with the atomistic MD simulation models, except for the GABARAP RKR region where we observed membrane association. One potential explanation is that the MD simulations model a single lipidated LC3B/GABARAP on a POPC membrane, whereas in the liposome-based assays, there might be multiple ATG8 proteins conjugated to one liposome, resulting conformational rearrangement of lipidated GABARAP (Coyle et al., 2002; Wu et al., 2015).

### Membrane insertion of LC3B/GABARAP N-termini is hindered by alteration of residues in loop 3 and RRR/RKR regions

As described above, we found that not only the N-termini, but also the loop 3 of LC3B/GABARAP associate with membranes in *cis* and that basic residues (RRR/RKR) are relatively detached from membranes after lipidation reaction. To further characterise the functions of the loop 3 and basic residues (RRR/RKR) in LC3B and GABARAP, we considered the apparent hydrophobicity of the residues in loop 3 (LC3B residues 42-44: KQL and GABARAP residues 39-41: ARI) and the RRR/RKR region and introduced triple glutamates (3E) which overall were either less hydrophobic or charge-inverting amino acids (Wimley and White, 1996). We employed N-terminal NBD labelled LC3B or GABARAP as a probe and assessed the real-time lipidation activity of these mutants (**Figure 4A**). No increases of NBD fluorescence were detected with LC3B S3C^NBD^ KQL-3E (loop 3 mutant), while lipidation still weakly occurred (**Figure 4B**). Moreover, for LC3B S3C^NBD^ RRR-3E (RRR mutant), the conjugation reaction was completely blocked (**Figure 4C**). Thus, mutations in LC3B loop 3 and RRR regions hindered lipidation due to the impaired conjugation reaction with ATG7/ATG3 and/or N-terminal membrane association.

**Figure 4.**
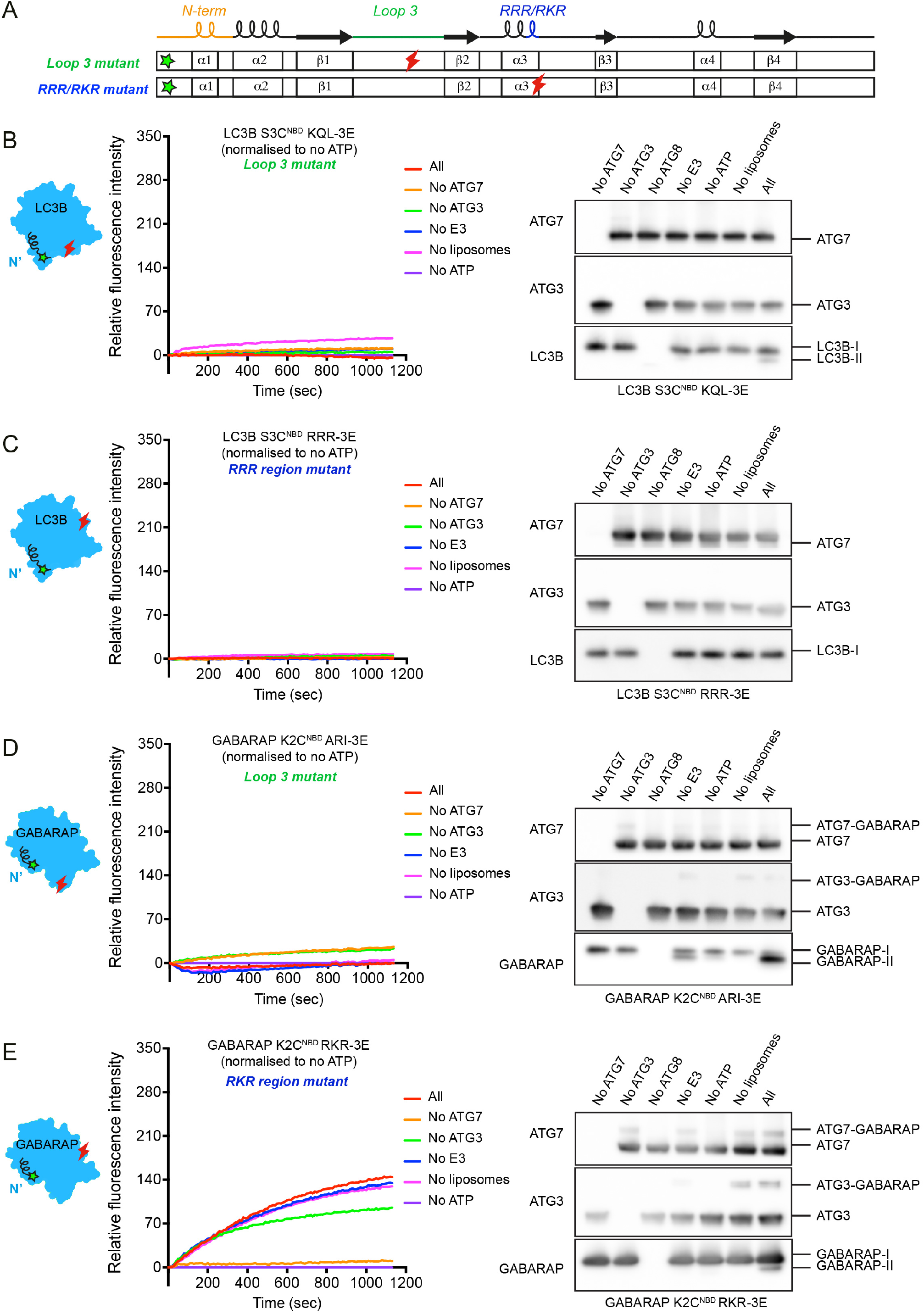
Membrane insertion of LC3B/GABARAP N-termini is hindered by alteration of residues in loop 3 and RRR/RKR regions. (**A**) Scheme of N-terminal NBD labelled LC3B (S3C^NBD^) or GABARAP (K2C^NBD^) with less hydrophobic and charge-inverting mutations (3E) in loop 3 and RRR/RKR regions. The green and red marks indicate the position of NBD labelling and 3E mutations, respectively. **(B)** The real-time lipidation assay with LC3B S3C^NBD^ KQL-3E (loop 3 mutant). **(C)** The real-time lipidation assay with LC3B S3C^NBD^ RRR-3E (RRR region mutant). (**D**) The real-time lipidation assay with GABARAP K2C^NBD^ ARI-3E (loop 3 mutant). (**E**) The real-time lipidation assay with GABARAP K2C^NBD^ RKR-3E (RKR region mutant). All spectra represent mean value (n=3).

Distinct from LC3B loop 3 mutant, GABARAP K2C^NBD^ loop 3 mutant (ARI-3E) was lipidated, while there was no increase in NBD fluorescence intensity (**Figure 4D**). The intermediates of GABARAP K2C^NBD^ ARI-3E mutant with ATG7 and ATG3 can also be observed although to a lower level compared to GABARAP K2C^NBD^ (**Figure 1E**). Thus, mutations in GABARAP loop 3 affected the conformation and hydrophobic environment of N-terminus during the lipidation reaction. With GABARAP K2C^NBD^ RKR-3E, the NBD fluorescence intensities were gradually increased. Different from GABARAP K2C^NBD^, the increased NBD signals indicated accumulations of ATG7∼/ATG3∼GABARAP intermediates (**Figure 4E**). GABARAP RKR mutant could not be discharged from ATG3∼GABARAP intermediate, resulting in impaired GABARAP-PE conjugation.

### Membrane insertion of LC3B/GABARAP N-termini contributes to efficient degradation of p62 body

We next investigated the effect of impaired membrane insertion of LC3B/GABARAP N-termini on autophagy in Hexa KO cells lacking all LC3/GABARAP subfamilies (Nguyen et al., 2016). We generated Hexa KO cell lines stably expressing GFP-fused GABARAP WT, an N-terminal deletion mutant (GABARAP Δ9), or impaired membrane association mutants (ARI-3E, RKR-3E). To minimize artificial effects of over-expression, each cell line was selected by FACS sorting for low levels of GFP expression. The cells were cultured under fed or starved conditions with and without lysosomal inhibitor bafilomycin A_1_ (BafA1) to check lysosomal degradation of autophagic cargos, GFP-GABARAPs and p62/SQSTM1, upon amino acid starvation. Starvation induced the degradation of GABARAP WT, GABARAP Δ9 and ARI-3E, while RKR-3E was not significantly degraded upon starvation (**Figure 5A and 5B**). These results indicate that GABARAP WT, GABARAP Δ9 and ARI-3E are efficiently incorporated into autophagic membranes and degraded by the autophagy-lysosome pathway. On the other hand, p62 degradation upon starvation was clearly observed only in the cells expressing GFP-GABARAP WT (**Figure 5A and 5C**). Hexa KO cells showed an accumulation of high molecular weight forms (high-MW) of p62 at approximately 170 kDa, which represents covalently crosslinked p62 oligomers (Donohue et al., 2014). This high-MW p62 was efficiently degraded in the cells expressing GFP-GABARAP WT as well (**Figure 5A and 5D**). Furthermore, S349-phosphorylated p62, which reflects the gel-like state of p62 (Kageyama et al., 2021), was clearly detected in Hexa KO cells, but completely suppressed by GFP-GABARAP WT expression (**Figure 5A and 5E**). In contrast, all GABARAP mutants did not significantly reduced levels of p62 and high-MW of p62 compared to GABARAP WT (**Figure 5A-5D**), indicating that p62 degradation can be affected by impaired *cis*-membrane association of LC3B/GABARAP N-termini even when the autophagy-lysosome pathway is still operational.

**Figure 5.**
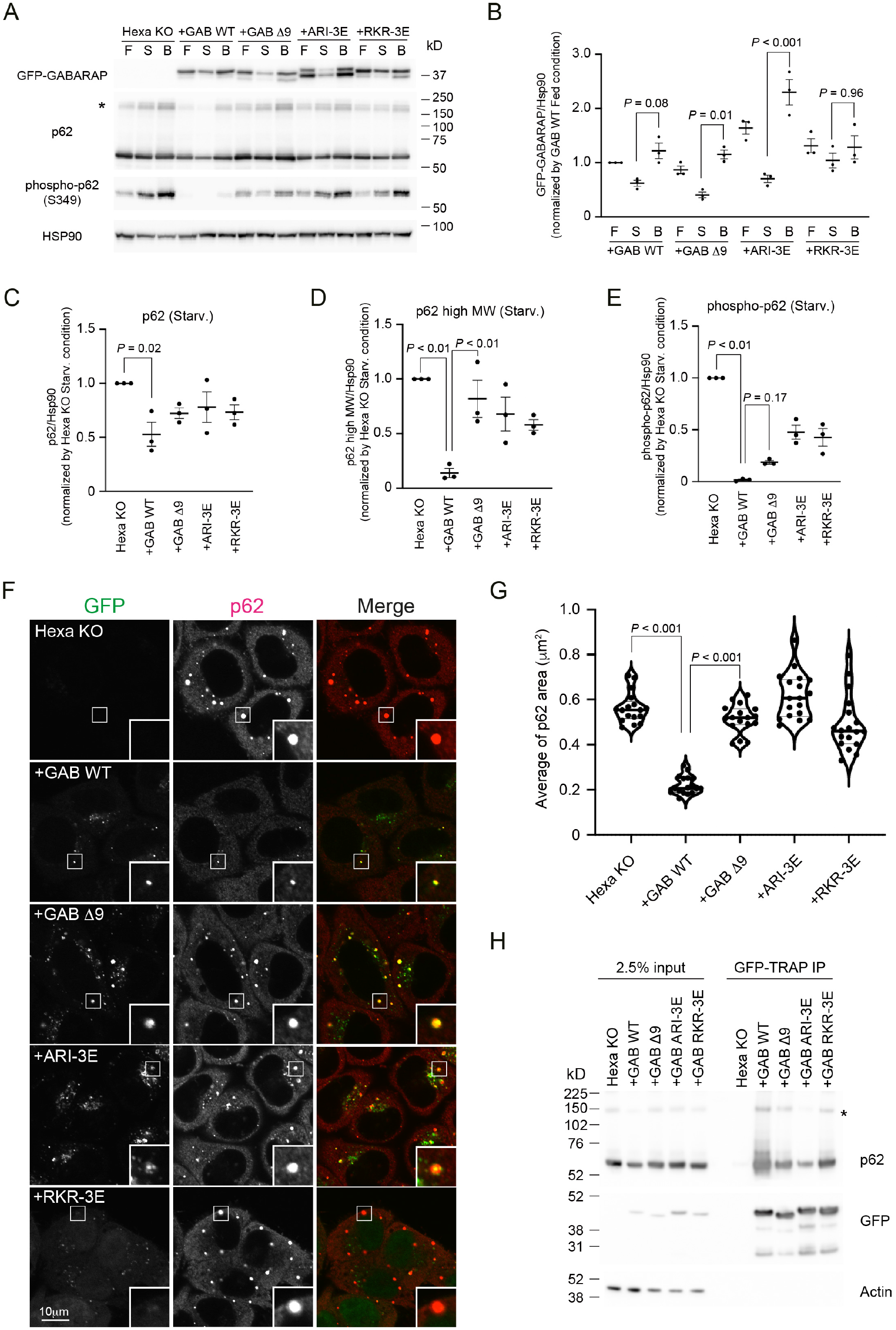
Membrane insertion of GABARAP N-terminus contributes to efficient degradation of p62 body. Hexa KO cells stably expressing GFP-GABARAP WT, Δ9, ARI-3E or RKR-3E were starved for 8 h with or without 100 nM Bafilomycin A_1_ (S), or cultured in full media (F). Cell lysates were analysed by immunoblotting using the indicated antibodies. The asterisk indicates the position of high molecular weight forms of p62. (**B-E**) Band intensity quantification of GFP-GABARAPs **(B)**, p62 **(C)**, p62 high molecular weight forms **(D)**, and phosphorylated p62 (Ser349) **(E)**. All data were normalized with those of HSP90. Data represent the mean ± SEM of three independent experiments. **(F)** The cells were starved for 2 h before p62 (red) was visualized. Scale bar, 10 μm. **(G)** The violin plot of average p62 area. The thick and thin lines in the violin plot represents the medians and quartiles, respectively. n = 17 for Hexa KO and GAB WT; n = 18 for GAB Δ9, ARI-3E or RKR-3E. **(H)** Co-immunoprecipitation of GFP-GABARAPs with p62. Cell lysates were subjected to IP using GFP-trap. The resulting precipitates were examined by immunoblot analysis with the indicated antibodies. Differences were statistically analysed by one-way ANOVA and Turkey multiple comparison test.

To gain further information, we further analysed the cellular localization of GFP-GABARAPs and p62 in the cells under starved conditions. In Hexa KO cells, p62 accumulated and formed large punctate structures (**Figure 5F**), which reflects impaired p62 degradation. Consistent with the observed rescue of degradation, GFP-GABARAP WT expression significantly reduced the size of p62 bodies (**Figure 5F**). The average area of the p62 bodies in cells expressing GFP-GABARAP WT was less than 40% of those observed in Hexa KO cells (**Figure 5G**). In contrast, the expression of the other GABARAP mutants did not cause a reduction in p62-body size (**Figure 5F and 5G**). Importantly, GFP-GABARAP mutants were still recruited to p62, forming small punctate structures on the surface of p62 bodies (**Figure 5F**). In line with this, co-immunoprecipitation experiments showed that GFP-GABARAP mutants retain the ability to interact with p62, though less efficiently compared to GFP-GABARAP WT (**Figure 5H**). Similar experiments were performed with LC3B where the reduction of p62 and high-MW of p62 were detected but less efficient (**Figure 5-figure supplement 1A-1E**), while GFP-LC3B WT efficiently reduced the levels of S349-phosphorylated p62. Quantification of the average of p62 bodies upon expression of GFP-LC3B N-terminal deletion mutant (Δ11) and the impaired membrane association mutants (KQL-3E and RRR-3E) confirmed that p62 bodies were not significantly reduced in size (**Figure 5-figure supplement 1F-1G**). Collectively, these results suggest that membrane insertion of LC3B and GABARAP N-termini is required for efficient degradation of p62 in cells.

### Membrane insertion of GABARAP N-terminus promotes phagophore expansion

As the GFP-GABARAP N-terminal deletion mutant (GABARAP Δ9) and impaired membrane association mutant (ARI-3E) partially co-localized with p62 puncta under starved condition (**Figure 5F**), we characterised the ultrastructure of these GFP-GABARAP-positive compartments attached to the p62 bodies, by correlative light-electron microscopy with serial EM sections (3D-CLEM). In Hexa KO cells, p62 bodies correlated with electron-dense structures (**Figure 6A**). Consecutive 25 nm SEM slices in the z-axis revealed that the size of the p62 body was more than 500 nm (**Figure 6B**, asterisks). In Hexa KO cells stably expressing GFP-GABARAP WT, the size of the p62 bodies was significantly reduced (**Figure 5G**) and no large electron-dense structures were observed (**Figure 6C**). Instead, GFP-GABARAP signals were correlated with double-membrane autophagosomes and multivesicular bodies (**Figure 6D**, arrowheads). In cells expression GFP-GABARAP Δ9 or GFP-GABARAP ARI-3E, the large p62 bodies were consistently correlated with electron-dense structures (**Figure 6E-H**, asterisks). Interestingly, 3D-CLEM analysis of these cells revealed that small autophagosome-like membrane were present and attached to the electron-dense structures in the area where GFP-GABARAP mutants and p62 were partially co-localized (**Figure 6F and 6H**, arrowheads). These results implicate the GABARAP N-terminus in the size regulation of autophagosomes and phagophore expansion.

**Figure 6.**
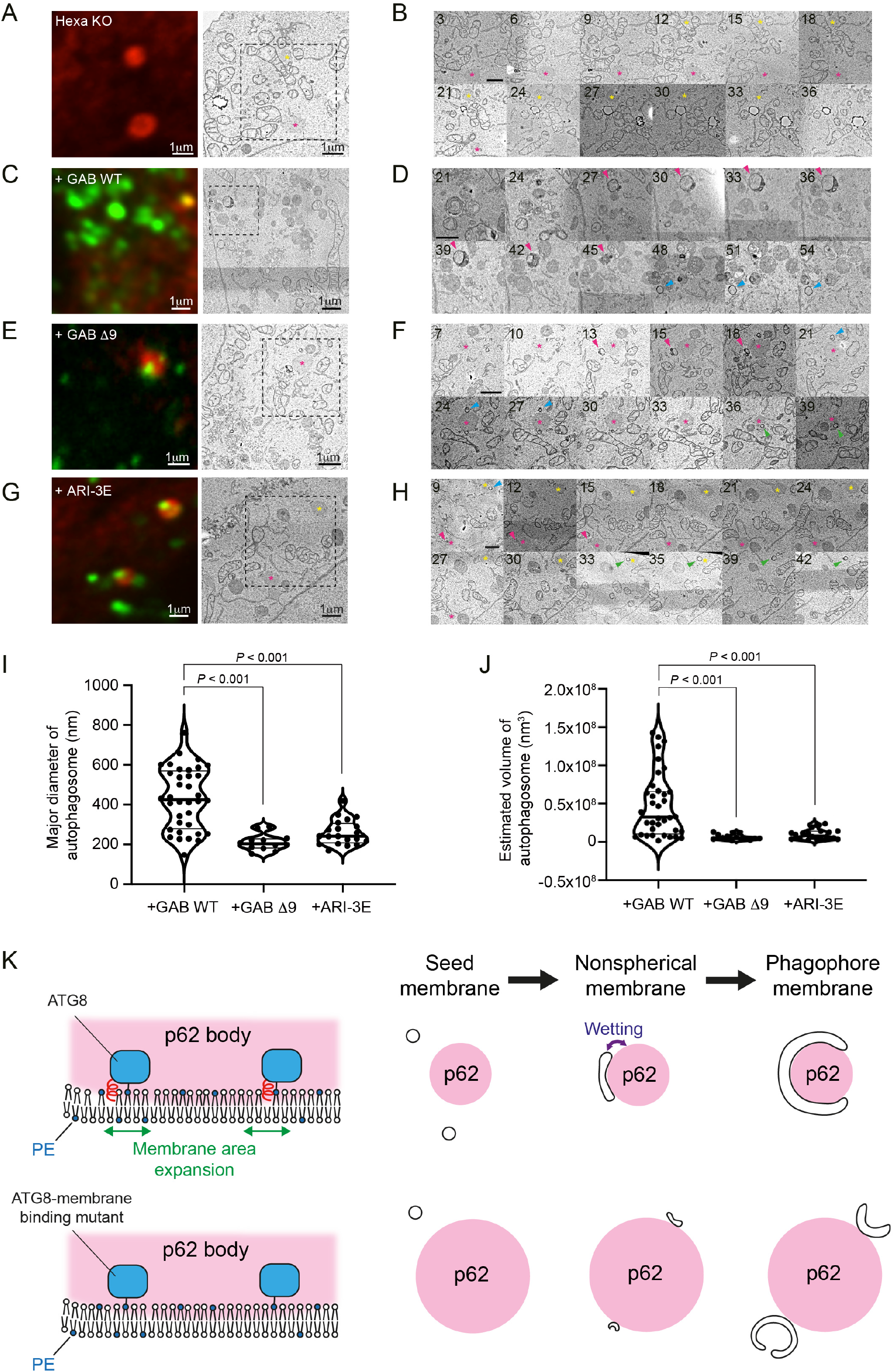
Membrane insertion of GABARAP N-termini promotes phagophore expansion. **(A-H)** Three-dimensional correlative light and electron microscopy (3D-CLEM) analysis. The starved cells were fixed, stained with p62, imaged by confocal microscopy, and subsequently relocated and imaged by scanning electron microscopy. Consecutive 25 nm SEM slice images of the area indicated by dushed boxes are shown (**B, D, F, H)**. The number in each image indicates the slice number. The asterisks and arrowheads show p62 bodies and autophagosomal structures, respectively. Scale bars, 1 μm. **(I, J)** Size distributions of autophagosomes. Length, width and height of each autophagosome are measured. The major diameter **(I)** and estimated volume **(J)** of each autophagosome are shown. The thick and thin lines in the violin plot represents the medians and quartiles, respectively. n = 36 for GAB WT; n = 12 for GAB Δ9; n = 20 for ARI-3E. **(K)** Model of autophagic membrane expansion and cargo degradation mediated by mammalian ATG8 proteins. After ATG8 lipidation, their N-termini are inserted into autophagic membranes. The cis-membrane association of the N-termini promotes autophagic membrane expansion by inducing membrane area expansion and helps efficient p62 degradation by facilitating wetting process which directs phagophore membranes toward p62 body.

To validate the possibility, we analysed the size of the autophagosomal structures correlated with GFP-GABARAP signals in a region of interest (ROI) of the 3D-CLEM data and estimated the volume of autophagosomes in each cell line (Maeda et al., 2020). The median of the major diameters of autophagosomal structures in Hexa KO cells expressing GABARAP WT was 426.3 nm, whereas those values significantly were reduced to 203.4 nm and 242.2 nm in the cells expressing GABARAP Δ9 and ARI-3E mutants, respectively (**Figure 6I**). In these cell lines, the estimated volume autophagosomes were less than 20% of those observed in the cells expressing GABARAP WT (**Figure 6J**). Taken together, these results support that the GABARAP N-terminus promotes phagophore expansion and increases the autophagosome sizes through its insertion into membranes.

Although we have not addressed this in our 3D-CLEM experiments, similar small punctate structures were observed attached to p62 in the LC3B mutants (Δ11 and KQL-3E) (**Figure 5-figure supplement 1F**), which are likely to be autophagosome-like structures and have a similar appearance as the GABARAP mutants (Δ9 and ARI-3E). This further supports the model we propose that the ATG8 N-termini promotes phagophore membrane expansion.

## Discussion

In this study, we investigated the dynamics of mammalian ATG8 N-termini during lipidation (**Figure 1**) and suggested a distinct conformation of lipidated ATG8 proteins, in which their N-termini are inserted into membrane *in cis* (**Figure 2 and 3**). The rescue assay using Hexa KO cells with ATG8 WT or ATG8 membrane-expansion mutants showed that the *cis*-membrane association of ATG8 proteins are critical for cargo degradation and autophagosome size regulation (**Figure 5 and 6**). Based on this evidence, we propose that the N-terminal regions of ATG8 proteins play critical roles in autophagy by associating with and being inserted into autophagic membranes (**Figure 6K**).

It was previously proposed that phagophore expansion is mediated by membrane tethering and fusion driven by ATG8 proteins anchored on two distinct membranes (“*in trans*”) (Nakatogawa et al., 2007; Weidberg et al., 2011; Wu et al., 2015). In this *trans* model, the N-terminus of lipidated Atg8 is expected to be solvent exposed. In support of this model, a recent study in yeast has resolved and modelled the structure of lipidated Atg8 on nanodiscs, revealing that two aromatic residues (F77/F79) of Atg8 are inserted into the membrane where lipidated Atg8 is present, thereby facilitating membrane expansion and deformation (Maruyama et al., 2021). In this model, the Atg8 N-terminus faces the cytoplasm, in contrast to our model of *cis*-membrane association of ATG8 N-terminus (that is, into the membrane where lipidated ATG8 is present). This discrepancy can be explained if ATG8 adopts distinct conformational states on autophagic membranes. Indeed, it has been reported that ATG8 proteins have open and closed conformations under non-lipidated state (Coyle et al., 2002; Wu et al., 2015). Given that ATG8 proteins are widely distributed on autophagic membranes (Sakai et al., 2020) and present during the progression from early to late stages of autophagosome formation, lipidated ATG8 proteins may undergo substantial conformational changes depending on their localization and the stages of phagophore growth. The convex and concave shapes of the phagophore membranes might be maintained by distinct conformation states of ATG8 proteins. Likewise, changes in membrane curvatures during autophagosome formation might have an impact on ATG8 conformation or even the membrane lipids themselves could be key players, particularly, PE. The ATG8-dependent membrane fusion and hemifusion activities require a high PE concentration, which is unusual in cell membranes (Nair et al., 2011). Our data show that lipid packing defects increased membrane association of ATG8 N-termini (**Figure 3D**). Accordingly, membrane environment may determine ATG8 conformation. How the structural dynamics of lipidated ATG8 is mediated by membrane curvature and membrane environment during autophagosome formation will be interesting to probe in the future research.

How does membrane insertion of ATG8 N-terminus induce phagophore expansion? One possible explanation is that membrane area is increased by space occupation and membrane deformation (Bangham, 1972; Maruyama et al., 2021). As ATG8 proteins are abundant on autophagic membranes, insertion of their N-termini into the membrane could have a large impact on autophagosome size. Another scenario is that local membrane elasticity on each side of the phagophore membrane might be regulated by ATG8 N-terminus, which would help to facilitate membrane expansion (Sakai et al., 2020). A recent paper reported that GABARAPs, but not LC3s, interact with the lipid transfer proteins ATG2A and B (Bozic et al., 2020). Therefore, GABARAP may have an additional role in mediating and combining ATG2-dependent lipid transfer with phagophore expansion.

The interaction of cargo receptors, such as p62 and NBR1, with LC3B/GABARAP directs phagophore membranes toward the cargos (Kageyama et al., 2021). This event, called wetting process, is crucial for efficient incorporation of cargos into autophagosomes (Agudo-Canalejo et al., 2021). We showed that impaired *cis*-membrane association mutants of GABARAP partially co-localized with p62 and did not fully restore degradation of the p62 bodies upon starvation in Hexa KO cells (**Figure 4 and 5**). However, we still observed the interaction of these GABARAP mutants with p62 in co-IP experiments, consistent with a previous report (Shvets et al., 2011). Therefore, abnormalities in p62 degradation do not simply result from a decreased ATG8-cargo receptor interaction. We interpreted these findings to show that in addition to ATG8-cargo receptor interaction, *cis*-membrane association of ATG8 proteins contributes to the wetting process. These two types of interactions may be additive and fine-tune membrane elongation along cargos. In the case of LC3B, it is hard to distinguish the function of the N-terminus in *cis*-membrane association from its interaction with p62, because the N-terminus is critical for the interaction with p62 (Shvets et al., 2008) and required for both events.

In this study, we newly developed an NBD fluorescence assay to monitor ATG8 lipidation reaction *in vitro* (**Figure 1**). The conjugation and lipidation cascades can be tracked by the incremental increase in the hydrophobic environment encountered by the N-termini of ATG8 proteins. These results clearly elucidated the highly dynamic nature of ATG8 N-termini during the lipidation reaction. Compared to giant unilamellar vesicles (GUVs)-based assay, our NBD lipidation assay is capable of measuring the ATG8 lipidation reaction on ∼100 nm LUVs in real-time. Therefore, the ATG8-PE conjugation can be investigated with membrane at a size more approximate to physiological size. Excitingly, these approaches can be extended to further studies of the effects of upstream factors, for example, the role of ATG2-dependent lipid transport (Maeda et al., 2019; Osawa et al., 2019; Valverde et al., 2019) on ATG8 lipidation.

The function of ATG8 proteins has been a long-standing question in autophagy research field. In this study, we have discovered the dynamic nature of ATG8 N-terminus and revealed how lipidated ATG8 regulates membrane expansion and cargo degradation. It should be noted that the *cis*-membrane association of ATG8 protein requires lipidation. Our findings have broader implications on the molecular mechanism of autophagosome formation and membrane growth, explaining why phagophore expansion requires ATG8 lipidation.

## Materials and Methods

### Plasmids

Descriptions of plasmids used in this study is in **Table S1**. For *in vitro* lipidation assay, pAL-GST-LC3B and pAL-GST-GABARAP were truncated to expose the C-terminal glycine (LC3B G120 and GABARAP G116). GABARAP-His_6_ construct was generated by introducing a His_6_-tag was after G116 in pAL-GST-GABARAP construct. LC3B-His_6_ construct was generated by amplifying DNA sequence encoding LC3B (aa 1-120) and subcloning the sequence into pGEX-6P1 vector with a His_6_-tag. All the point mutations in the recombinant LC3B and GABARAP were generated by Q5 Site-Directed Mutagenesis Kit (New England BioLabs). pGEX-6P1-GST-ATG3 was a gift from Dr. Alicia Alonso. cDNA encoding full-length human ATG7 (provided by Dr. Alicia Alonso, Instituto Biofisika, University of the Basque Country) was amplified by PCR and subcloned into pBacPAK-His_3_-GST construct (provided by Dr. Svend Kjaer, The Francis Crick Institute) via In-Fusion cloning method (Takara Bio). StrepII^2x^-ATG5, ATG7, ATG10, ATG12 and ATG16L1 were cloned into a single pFBDM plasmid.

For generation of stable retroviral transduced cell lines, cDNAs encoding LC3B (NM_022818) and GABARAP (NM_007278) were amplified by PCR and subcloned into pMRX-IP backbone vector together with the EGFP tag by Gibson Assembly (New England Biolabs, E2611L). All deletion and point mutation mutants of LC3B and GABARAP were generated by a PCR-based method.

### Antibodies and reagents

Antibodies used for immunoblotting are listed: rabbit polyclonal anti-ATG7 (Cell Signalling Technology, 2631), rabbit polyclonal anti-ATG3 (Sigma, A3231), rabbit polyclonal anti-LC3B (Abcam, ab48394), rabbit monoclonal anti-GABARAP (Cell Signalling Technology, 13733), mouse monoclonal anti-GABARAP (MBL, M135-3) (for blotting GABARAP K2C^NBD^ ARI-3E mutant only), mouse monoclonal anti-HSP90 (BD Transduction Laboratories, 610419), rabbit polyclonal anti-p62 (MBL, PM045, lot 022), rabbit polyclonal anti-Phospho-p62 Ser351 (MBL, PM074), rabbit polyclonal anti-GFP (Invitrogen, A6455), mouse monoclonal anti-GFP (Cancer Research UK (raised in-house)), rabbit polyclonal anti-Actin (Abcam, ab8227). Secondary antibodies are HRP-conjugated anti-rabbit IgG (Jackson ImmunoResearch Laboratories, 111-035-144 or GE Healthcare, #NA934), HRP-conjugated anti-mouse IgG (Jackson ImmunoResearch Laboratories, 315-035-003 or GE Healthcare, #NA931).

Antibodies used for immunofluorescence are as listed: rabbit polyclonal anti-p62 (MBL, PM045, lot 022) and AlexaFluor 568-conjugated anti-rabbit IgG (Invitrogen, A-11036).

All lipids were purchased from Avanti: POPC (850457 C), DOPC (850375 C), POPE (850757 C), DOPE (850725 C), DGS-Ni-NTA (790404C), Rhod-PE (810150 C). ATP solution (100mM) was obtained from ThermoFisher (R0441). IANBD was purchased from Invitrogen (D2004).

### Recombinant protein expression and purification

ATG3, LC3B, GABARAP and their corresponding mutants were transformed in *Eschericia coli* BL21 (DE3) cells. Bacteria were grown in LB until OD_600_ = 0.8 and protein expression was induced with 0.5mM IPTG for 16hr at 18°C with all the constructs, except that in the case of GABARAP K2C ARI-3E, bacteria were grown in TB and protein expression was induced when OD_600_=1.2. These GST-tagged proteins were purified as described before with minor modifications (Wirth et al., 2019). Briefly, cells were harvested by centrifugation and resuspended in 50 mM Tris-HCl pH8.0, 500 mM NaCl, 0.5 mM TCEP, 0.4 mM AEBSF, and 15 µg/ml benzamidine. Cells were lysed by freeze-thaw followed by sonication. Lysates were then cleared by centrifugation at 25,000g for 30min at 4°C. The GST-tagged proteins were absorbed with Glutathione-Sepharose 4B affinity matrix (GE Healthcare) for 1.5hrs and recovered by 3C protease cleavage at 4°C overnight in 50 mM Tris-HCl pH 8.0, 500 mM NaCl and 0.5 mM TCEP. The proteins were further purified by size exclusion chromatography using Superdex 200 16/60 column (GE Healthcare) equilibrated in buffer containing 25mM Tris-HCl pH8.0, 150mM NaCl and 0.5mM TCEP.

To purify human ATG7, pBacPAK-His_3_-GST-ATG7 was transfected into insect cells Sf9 using the FlashBAC baculovirus expression system (Oxford Expression Technologies OET) and Fugene HD transfection reagent (Promega) according to manufacturer’s instructions. Virus at P1 was harvested after 5 days from transfection. 50mL of Sf9 cells were infected with 1.5mL of P1 virus to amplify the virus and harvest the stock virus P2. 200mL of Sf9 cells (1.5-2×10^6^ cells/mL) was infected with 1mL of P2 virus and harvested after 60hrs. The cell pellets were frozen in liquid nitrogen and stored at -80°C until purification. When purifying ATG7, the cells were thawed on ice and resuspended in lysis buffer containing 50 mM Tris-HCl pH 8.0, 500 mM NaCl, 0.5 mM TCEP and EDTA-free Complete Protease inhibitor cocktail (Roche). The suspension was sonicated 10sec, 5 times on ice, and then centrifuged at 30,000g for 30min at 4°C to remove cell debris. The remaining purification steps were performed as described above.

To purify human E3 complex, pFBDM-ATG7-ATG10-ATG12-StrepII^2x^-ATG5-ATG16L1 was transformed into DH10Multibac cells. The bacmid was isolated and Sf9 cells transfected. The resultant baculovirus was further amplified using standard procedures. The ATG12–ATG5-ATG16L1 complex was expressed in High Five insect cells using Sf-900 II SFM medium (Thermo). Cells were infected with a multiplicity of infection (MOI) greater than 2. Cells were harvested after 2.5 days and lysed using pre-cooled lysis buffer containing 50 mM Tris HCl pH 8.3, 220 mM NaCl, 5% glycerol, 2 mM DTT, EDTA-free protease inhibitor tablets (Roche), 2 mM EDTA, 0.2 mM PMSF, 1 mM benzamidine and Pierce universal nuclease. Cells were lysed by sonication and spun at 20,000 rpm for one hour using a JA-20 rotor. The supernatant was loaded onto a StrepTactin column (Qiagen) pre-equilibrated with wash buffer composed of 50 mM Tris HCl pH 8.0, 220 mM NaCl, 5% glycerol and 2 mM DTT. The StrepTactin column was washed with 20 column volumes (CV) of wash buffer before the E3 complex was eluted with 5 CV wash buffer containing 2.5 mM desthiobiotin. E3 containing fractions were pooled and loaded onto a ResQ anion exchange chromatography column (GE Healthcare). The E3 complex was eluted by applying a salt gradient from 50 to 700 mM NaCl (ResQ buffer base: 20 mM HEPES-NaOH pH 8.0, 5% glycerol and 2 mM DTT). Protein containing fractions were pooled, concentrated and loaded on a HiLoad 16/600 Superose 6 size exclusion chromatography column pre-equilibrated in SEC buffer (20 mM HEPES NaOH pH 7.4, 180 mM NaCl, 5% glycerol and 2 mM DTT). The E3 complex was concentrated using Amicon Ultra concentrators, aliquoted and flash-frozen in liquid nitrogen.

### Liposome preparations

Lipids were mixed at the desired molar ratio in chloroform, dried under nitrogen gas and further vacuumed for 2hrs to remove the remaining solvent. The lipid film was rehydrated and resuspended in the assay buffer containing 25mM Tris-HCl pH 8.0, 150mM NaCl and 0.5mM TCEP, with vortexing. Large unilameller vesicles (LUVs) were generated by 5 times freeze-thaw cycles in liquid nitrogen and water bath. The LUVs then were extruded 10 times through 0.2μm membrane followed by at least 20 times through 0.1μm membrane (Whatman) using a Mini-Extruder (Avanti Polar Lipid). The final concentration of liposome was 2mM. The size of LUVs was checked by Zetasizer Nano ZS (Malvern Instruments). The LUVs had an average diameter of 100nm. All the *in vitro* fluorescence measurements (the real-time lipidation assay, fluorescence-based liposome binding assay and FRET assay) were performed in the same assay buffer used for liposome preparation.

### Protein NBD labelling

NBD is an environment-sensitive fluorescence probe with low molecular weight. It has been commonly used to monitor the environmental changes of specific residues of proteins (Raghuraman et al., 2019). LC3B and GABARAP proteins with a single cysteine mutation were labelled as described with some modifications (Nishimura et al., 2019). The purified proteins were transferred to labelling buffer Tris-HCl pH7.5 20mM, NaCl 150mM using a PD Minitrap G-10 column (Cytiva), immediately prior to labelling. The proteins were then labelled with 20-fold excess of IANBD-amide at room temperature in dark for 1hr. The reaction was terminated with 4mM cysteine and excess of IANBD was removed by a PD Minitrap G-10 column, preequilibrated with 20mM Tris-HCl pH 7.5, 150mM NaCl and 0.5mM TCEP. Glycerol was added to the NBD labelled protein to a final concentration of 20% for storage at -80°C. The concentration of NBD labelled proteins were determined by Bradford and checked by SDS-PAGE. The concentrations of the labelled proteins ranged between 30μM to 80μM.

### Real-time lipidation assay

All the real-time lipidation assays were carried out at 37°C with three independent repeats. Briefly, the reaction mix (80μL) except the ATP/MgCl_2_ was prepared with 0.2μM ATG7, 0.2μM ATG3, 1μM NBD-labelled LC3B/GABARAP, 0.05μM the E3 complex and 1mM liposomes. The reaction mix was then transferred into 10-mm pathlength quartz cuvette and the NBD fluorescence (ex/em 468nm/535nm) was measured immediately using FP-8300 spectrofluorometer (JASCO). The excitation bandwidth was fixed to 5nm, and the emission bandwidth was fixed to 10nm. The total measurement time was set to 20min with two time intervals: in the first 80sec, NBD fluorescence was recorded every 20sec; after 80sec, the fluorescence was recorded every 10sec until the end. ATP/MgCl_2_ (final conc. 1mM) was added into the reaction mix between 60sec and 80sec to initiate the lipidation reaction. For the control group containing all the proteins and liposomes except for the ATP/MgCl_2_, the same amount of buffer was added instead. The fluorescence increase (ΔEm535nm) at each time point was calculated by subtracting the NBD signal recorded from the control group, normalised to the time point at 80sec.

For step-by-step assay (**Figure 1F**), there were some modifications with data collection intervals. The total measurement time was set to 1320sec, and the NBD fluorescence was recorded every 20sec. There were four reaction groups: (1) hATG8^NBD^, (2) hATG8^NBD^ + liposomes, (3) hATG8^NBD^ + ATP/MgCl_2_ and (4) hATG8^NBD^ + liposomes + ATP/MgCl_2_. ATG7 (final conc. 0.2μM) was added between 60sec-80sec. ATG3 (final conc. 0.2μM) was added between 480sec-500sec. The E3 complex (final conc. 0.05μM) was added between 900sec-920sec. There were two control groups, (1) hATG8^NBD^ and (2) hATG8^NBD^ + liposomes. Instead of adding ATG7, ATG3 and the E3 complex, the same amount of buffer was added in between each corresponding time interval. The reaction (1) and (3) were calibrated to the control group (1), subtracting the background signal of hATG8^NBD^. The reaction (2) and (4) were calibrated to the control group (2), subtracting the background signal of hATG8^NBD^ and liposomes. We determined the fluorescence increase (ΔEm535nm) at each time point normalised to the fluorescence recorded at the time point after the addition of ATG7 (80sec), ATG3 (500sec) and the E3 complex (920sec).

### Fluorescence-based liposome binding assay and FRET assay

The spectra of fluorescence-based liposome binding assay and FRET assay were recorded at 37°C. For fluorescence-based liposome binding assay, NBD labelled LC3B-His_6_/GABARAP-His_6_ (1μM) were mixed with liposomes (final conc.1mM) in a total volume of 80μL and immediately measured using FP-8300 spectrofluorometer. The emission spectra were recorded from 500nm to 650nm by exciting NBD at 468nm. The excitation and emission bandwidths were set to 5nm. For FRET assays, the settings of fluorescence measurements were kept the same. The NBD-labelled LC3B-His_6_/GABARAP-His_6_ (1μM) were mixed with blank liposomes or rhodamine liposomes (final conc. 1mM), in the presence or absence of imidazole (final conc. 100mM). The NBD fluorescence was recorded immediately. The spectra of buffer and all liposomes solutions were recorded as background control. The spectra of NBD-labelled proteins were corrected by subtracting the buffer spectra, while the reactions containing liposomes were corrected by subtracting the corresponding liposome spectra.

### Molecular dynamic simulations

The protein structure of LC3B (3VTU) and GABARAP (1GNU) protein were used and residues after Gly-120 and Gly-116 were deleted (Jatana et al., 2020). A covalent bond between C-terminal Glycine and phosphatidylethanolamine (PE) was introduced where the lipid anchor parameters were taken from already existing 1-palmitoyl-2-oleoly-*sn*-phosphoethanolamine (POPE) of charmm36 ff. The lipidated LC3B/GABARAP proteins were placed in a 400-lipid POPC bilayer membrane which was prepared using charmm-GUI web server. To obtain statistically accurate docking of LC3 on membrane, we undertook multiple conformations with varied lipid-contacting orientations (**Figure 2**). After an initial refinement of 100 ns, all conformations converged to a single orientation with N-terminal moving closer to lipids. Hence, the conformation with membrane facing N-terminal was taken for further simulations. In total three simulations each were started for LC3 and GABARAP and cumulative simulation length was 6.6 μs was obtained.

The lipidated structures of LC3B/GABARAP with POPC membrane were placed in a rectangular box large enough to accommodate protein and membrane. Water molecules were added with TIP3P representation and Na^+^ ions were added to neutralize the systems (Mark and Nilsson, 2001). MD simulation was performed using GROMACS version 2018.3 by utilizing charmm36 all-atom FF (Abraham et al., 2015). Periodic boundary conditions were used, and 1.2 nm was set as real space cut-off distance. Particle Mesh Ewald (PME) summation using the grid spacing of 0.16 nm was used in combination with a fourth-order cubic interpolation to deduce the forces and potential in-between grid points (Darden et al., 1993). The Van der Waals cut-off was set to 1.2 nm. Energy minimization was performed on the initial systems using the steepest descent method and the temperature and pressure were maintained at 310 K and 1 bar using Nose-Hoover thermostat and Parrinello-Rahman barostat, respectively (Bussi et al., 2007; Parrinello and Rahman, 1981). A time step of 2 fs was used for numerical integration of the equation of motion. The coordinates were saved at every 20 ps. Three replicas for each lipidated LC3B and GABARAP systems were simulated for 1µs. All the molecular images were rendered using UCSF Chimera and VMD (Humphrey et al., 1996). The graphs and plots were generated using MATLAB and Python libraries.

### Analysis of trajectories

The periodic boundary conditions were removed before performing analysis on the trajectories. The distance was calculated between the centre of mass of residues and the POPC membrane was calculated using the gmx distance module which calculates the distance between two positions as a function of time. The probability of contact formation was calculated be defining a contact when the distance is between residues and membrane was <3.5 nm. Further the contacts were categorized into three parts depending upon the residues’ side chain orientation on the membrane: Inserted at ≤ 2.3nm, at membrane surface at ≤ 2.8nm, in proximity at < 3.5nm.

### Cell lines and culture conditions

Authenticated human embryonic kidney (HEK) 293T cells were used in this study. Hexa KO cell line was generated previously (Nguyen et al., 2016). Cells were maintained in Dulbecco’s Modified Eagle Medium (DMEM) (Wako, 043-30085) supplemented with 10% fetal bovine serum (FBS) (Sigma-Aldrich, 173012) in a 5% CO_2_ incubator at 37 °C. Hexa KO cells stably expressing GFP-fused GABARAPs were generated as follows: HEK293T cells were transfected using Lipofectamine 2000 reagent (Thermo Fisher Scientific, 11668019) with pMRX-IP-based retroviral plasmid, pCG-VSV-G and pCG-gag-pol, following which the medium was replaced with fresh medium. After 3 days, the culture medium was collected and filtered with a 0.45 μm filter unit (Millipore, SLHVR33RB). Hexa KO cells were treated with the retrovirus containing medium and 8 μg/mL polybrene (Sigma-Aldrich, H9268). After 2 days, drug selection was performed with 3 μg/mL puromycin (Sigma-Aldrich, P8833). Puromycin-resistant cells were passaged 3 times and then sorted by flow cytometry (Sony, SH800) to select GFP-low expressing cells.

### Immunoblotting

Cells were cultured under DMEM supplemented with FBS or DMEM without amino acids (Wako, 048-33575) in the absence or presence of 100 nM bafilomycin A_1_ (invitrogen, B1793) for 8 h, collected in ice-cold PBS by scraping and then precipitated by centrifuged at 1,000 x g for 3 min. The precipitated cells were suspended in 100 μl lysis buffer (25 mM Hepes-NaOH, pH7.5, 150 mM NaCl, 2 mM MgCl_2_, 0.2% *n*-dodecyl-b-D-maltoside [nacalai, 14239-54] and protease inhibitor cocktail [nacalai, 03969-34]) and incubated on ice for 20 min. Ninety μl of cell lysates were mixed with 10 μl of lysis buffer containing 0.1 μl benzonase (Merck Millipore, 70664) and further incubated on ice for 15 min. The remaining cell lysates were centrifuged at 17,700 x g for 15 min and the supernatant was used to measure protein concentration by NanoDrop One spectrophotometer (Thermo Fisher Scientific). One hundred μl of cell lysates were mixed with SDS-PAGE sample buffer and heated at 95°C for 5 min. Samples were subsequently separated by SDS-PAGE and transferred to Immobilon-P PVDF membranes (Merck Millipore, IPVH00010) with Trans-Blot Turbo Transfer System (Bio-Rad). After incubation with the indicated antibodies, the signals from incubation with SuperSignal West Pico PLUS Chemiluminescent Substrate (Thermo Fisher Scientific, 34580) was detected with Fusion Solo S (VILBER). Band intensities were quantified with Fiji.

For *in vitro* real-time lipidation reaction, 20μL reaction mix was taken immediately after the reaction, mixed with 5μL SDS-PAGE sample buffer and heated at 95°C for 5min. 10μL reaction mix was resolved on NuPAGE Bis-Tris 4–12% gels (Life Technologies) followed by immunoblotting.

### Immunoprecipitation

Cells were lysed in ice cold lysis buffer containing 20mM Tris-HCl 7.4, 150mM NaCl, 150mM EDTA, 0.5% w/v Triton X-100, 1× Complete protease inhibitor (Roche) and 1× PhosSTOP (Roche). Cell lysate was centrifuged at 17,700 x g for 15min. The supernatant was precleared with binding control agarose beads (ChromoTek) for 1hr at 4°C. GFP-GABARAP proteins were immunoprecipitated using GFP-TRAP beads (ChromoTek), incubating for 2hrs at 4°C. After washing beads 4 times with lysis buffer (w/o PhosSTOP), beads were mixed with SDS-PAGE sample buffer and boiled at 95°C for 5min. The proteins of interest were resolved on NuPAGE Bis-Tris 4–12% gels (Life Technologies) followed by western blotting.

### Fluorescence microscopy

Cells grown on coverslips were fixed with 4% paraformaldehyde in PBS for 15 min, permeabilized with 50 μg/ml digitonin (D141; Sigma-Aldrich) in PBS for 5 min, blocked with 3% BSA in PBS for 30 min, and then incubated with anti-p62 antibody for 1h. After washing five times with PBS, cells were incubated with Alexa Fluor 568 conjugated goat anti-rabbit IgG secondary antibody for 1h. These specimens were observed using a confocal FV3000 confocal laser microscope system (Olympus). For the final output, images were processed using Adobe Photoshop 2021 v22.3.1 software (Adobe). Average of p62 were measured using the open-source software Fiji.

### Correlative light and electron microscopy

Correlative light and electron microscopic (CLEM) analysis was performed as previously described (Maeda et al., 2020; Morishita et al., 2021). In brief, Hexa KO, Hexa KO stably expressing GFP-GABARAP, GFP-GABARAP Δ9, or GFP-GABARAP ARI-3E, were grown for two nights in custom made gridded coverslip-bottom dishes. The cells were fixed, stained with anti-p62 antibody and observed by FV3000 confocal laser microscope system (Olympus) equipped with a 60× oil-immersion objective lens (NA1.4, PLAPON60XOSC2; Olympus). After fluorescent-image acquisition, the cells were embedded in epoxy resin for electron microscopic observation as previously described (McArthur et al., 2018; Morishita et al., 2021). After embedding and removal of the coverslip, the resin block was trimmed to remain the imaged area by confocal laser scanning microscope (about 150 × 150 μm). Then the block was sectioned using ultramicrotome (EM UC7, Laica) equipped with a diamond knife (Ultra JUMBO, 45 degrees, DiATOME) to cut 25 nm serial sections. With an active vibration isolation table and an eliminator of the static electricity were used to make long serial sections. Then the sections were collected on the cleaned silicon wafer strip held by a micromanipulator (Märzhäuser Wetzlar). Those sections were directly imaged (without staining) using a scanning electron microscope (JSM7900, JEOL) supported by a software (Array Tomography Supporter, System in Frontier). Images were stacked in order by a software (Stacker NEO, System in Frontier). The correlation of light and electron microscopic images was performed by FIJI software or Adobe Photoshop. Before quantification of autophagosome sizes, the samples were shuffled for a randomized double-blind analysis. The diameters of autophagosomal structures in the maximum cross section were measured by FIJI software. The total number of slices showing autophagosomes was used to measure their heights. Autophagosome volume was calculated using the formula for volume of ellipsoid sphere: V = length x width x height x π/6.

### Data analysis

Differences were statistically analysed by one-way ANOVA and Turkey multiple comparison test. Statistical analysis was carried out using GraphPad Prism 9 (GraphPad Software).

## Data availability

All data generated or analysed during this study are included in the manuscript or supporting files. Spectra data, gels and blots are included in Source data file.

## Acknowledgements

We thank Noboru Mizushima (N.M.), the director of Exploratory Research for Advanced Technology (ERATO) Mizushima Intracellular Degradation project, for helpful discussion, Yoko Ishida for cutting ultrathin sections and providing electron microscopy pictures, Ikuko Koyama-Honda, Satoru Takahashi, Keiko Igarashi for technical assistance with the 3D-CLEM experiments, Shoji Yamaoka for pMRXIP, and Teruhito Yasui for pCG-VSV-G and pCG-gag-pol. We thank Michael Lazarou for providing Hexa KO cell line, Alicia Alonso for human ATG7 and ATG3 plasmids, Svend Kjaer for pBacPAK-His_3_-GST plasmid and the technical assistance with insect cell culture, Simone Kunzelmann for the technical assistance and helpful suggestions on fluorescence spectroscopy, and Stefano De Tito for critical comments and suggestions. We thank the support from CSIR-IGIB for infrastructure and CSIR-4PI for supercomputing facilities. This study was supported by PRESTO (JPMJPR20EC to T.N.) and ERATO (JPMJER1702 to C.S. via N.M.) from Japan Science and Technology (JST), a Grant-in-Aid for Transformative Research Areas (B) (grant 21H05146 to T.N.) from the Japan Society for the Promotion of Science (JSPS), and a grant from the Japan Foundation for Applied Enzymology (to T.N.). D.G. and L.T. were supported by funding from CSIR and the Department of Science and Technology, CSIR-NET for fellowship and (DST)-SERB Early Career Grant and OLP1163. W.Z., H.B.J.J, C.D., A.S. and S.A.T, were supported by The Francis Crick Institute which receives its core funding from Cancer Research UK (CC2134, CC2064), the UK Medical Research Council (CC2134, CC2064). This research was funded in whole, or in part, by the Wellcome Trust (CC2134, CC2064). W.Z. and S.A.T received funding from the European Research Council under the European Union’s Seventh Framework Programme (FP7/2007-2013)/ERC grant agreement n° [788708]. For the purpose of Open Access, the author has applied a CC BY public copyright licence to any Author Accepted Manuscript version arising from this submission.

## Author Contributions

W.Z. and T.N. performed the majority of the experimental work. T.N. designed the real time assay. D.G. and L.T. performed the simulation analysis. T.N. and C.S. performed the CLEM experiments and analysis. W.Z., T.N., C.D. and A.S. purified proteins. H.B.J.J. assisted with the stable cell generation. W.Z., T.N. and S.A.T. designed this study and wrote the manuscript with input from all authors.

---

**Figure 1-figure supplement 1.**
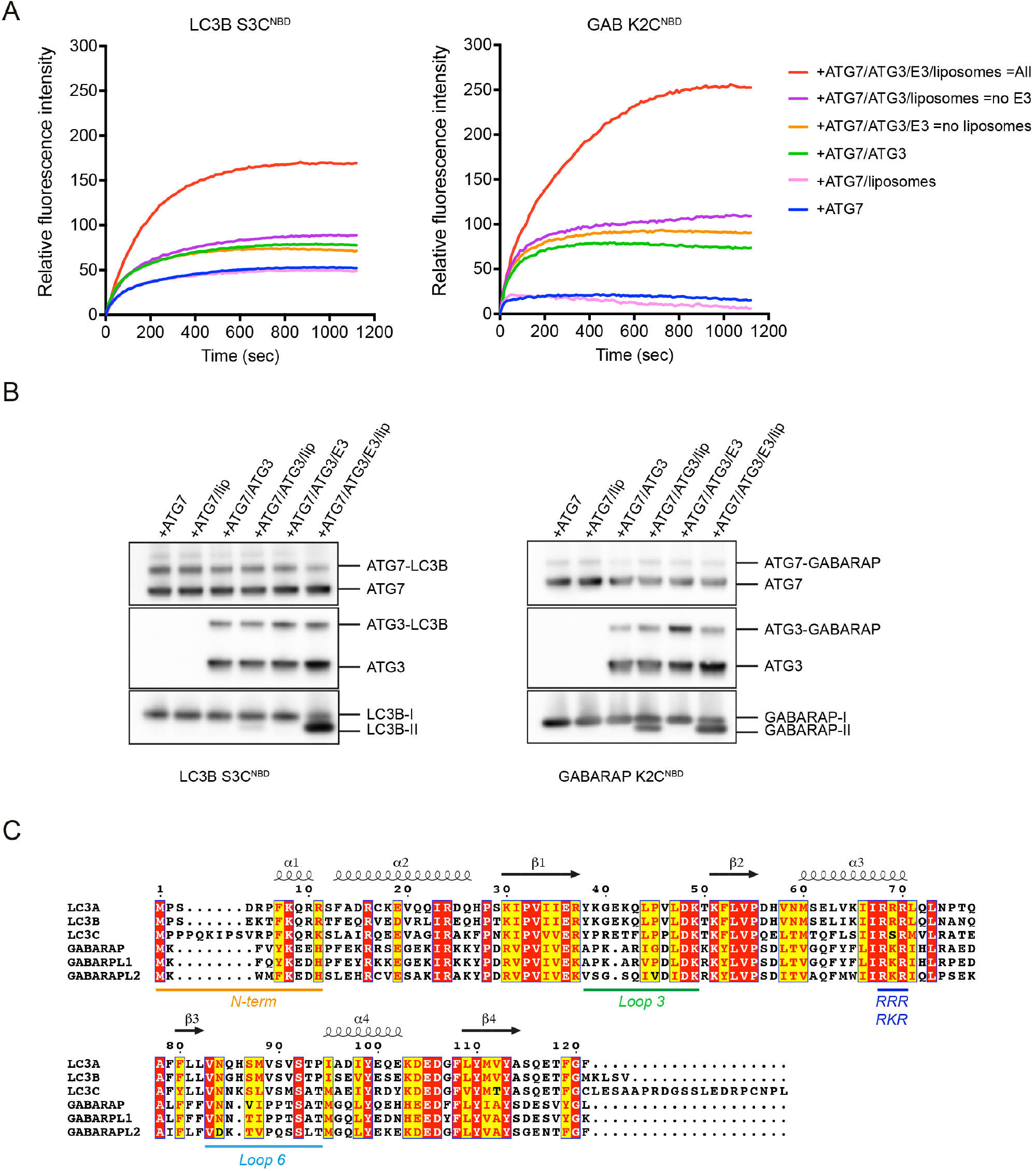
The increase of LC3B/GABARAP N-terminal NBD fluorescence reflects the conjugation with ATG7, ATG3 and lipidation of LC3B/GABARAP. (**A**) Control experiments for the real-time assay with LC3B S3C^NBD^ and GABARAP K2C^NBD^ in Fig. 1. Instead of removing one component from the “All” conditions, the assays were performed with components indicated in the legend. The NBD fluorescence was recorded once ATP was added (time point at 0s). The relative NBD fluorescence was normalised to that of LC3B S3C^NBD^ and GABARAP K2C^NBD^ protein only, respectively. (**B**) Western blots of sample from (**A**). (**C**) Sequence alignment of six human ATG8 proteins using ESPript 3. The identical and similar residues are indicated in red and yellow, respectively. N-terminus (orange), loop 3 (green), basic patch RRR/RKR (blue) and loop 6 (light blue) were shown.

**Figure 2-figure supplement 1.**
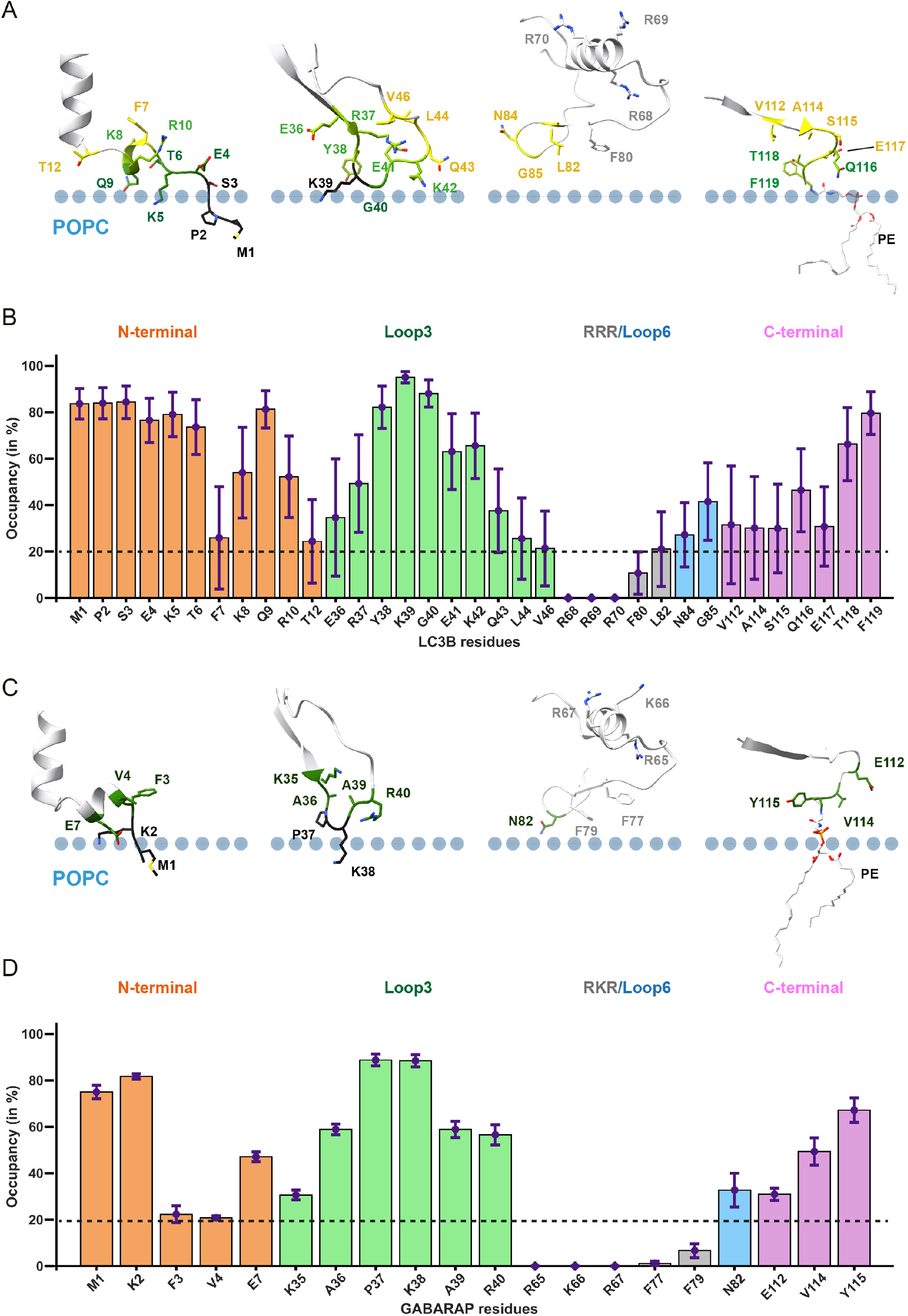
Side-chain orientation of membrane interacting residues. **(A) and (C)** Snapshots highlighting the side-chain orientation of membrane interacting regions mapped on the representative structure of LC3B (300ns) and GABARAP (840ns) respectively. The position of membrane is highlighted as dot-line. **(B) and (D)** Showing the percentage occupancy of specific residues. The bars are coloured according to the region which is N-terminal as (orange), loop 3 as green, loop 6 as light blue, and C-terminal as purple.

**Figure 3-figure supplement 1.**
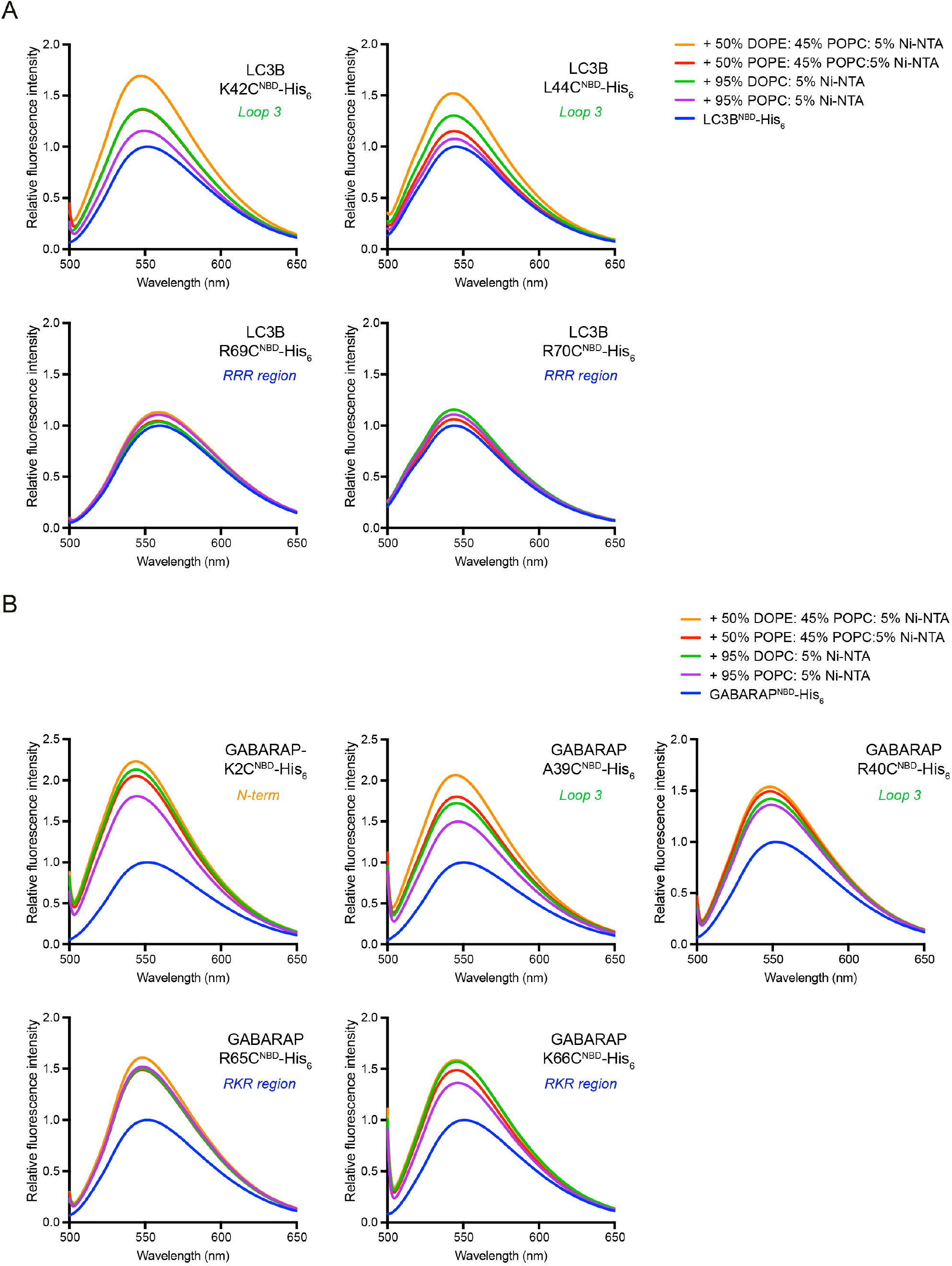
Relative fluorescence spectra of NBD-labelled and liposome conjugated LC3B/GABARAP. (**A**) LC3B loop 3 (green) and RRR region (blue) were tested for membrane binding with selected residues. (**B**) GABARAP N-terminal (orange), loop 3 (green) and RKR region (blue) were tested for membrane-association activity with selected residues. Spectra represent mean values (n=4-5)

**Figure 3-figure supplement 2.**
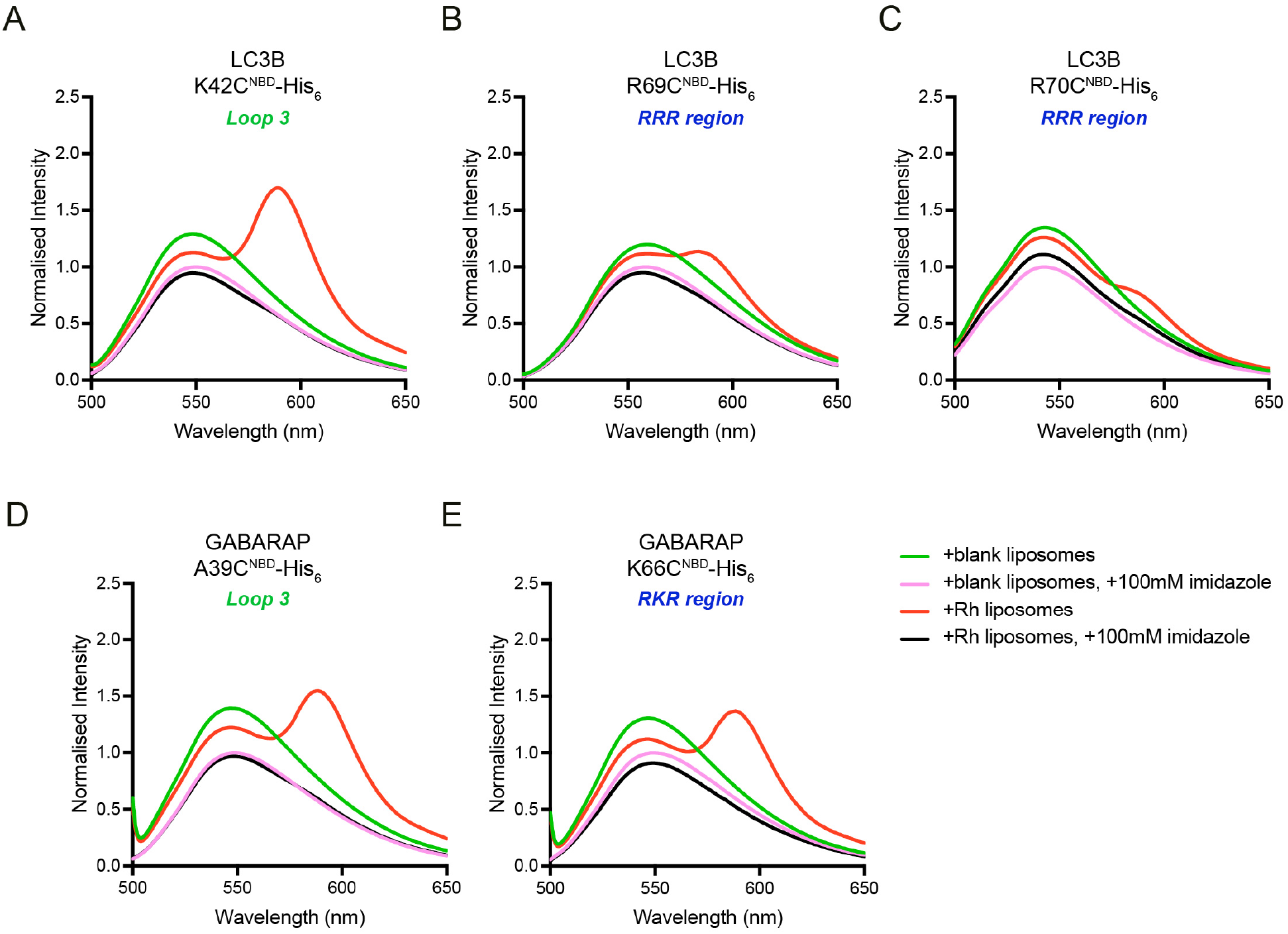
FRET between NBD-labelled LC3B-His_6_/GABARAP-His_6_ and rhodamine-labelled liposomes. **(A-C)** FRET assay with NBD labelled residues in LC3B-His_6_. **(D-E)** FRET assay with NBD labelled residues in GABARAP-His_6_. Spectra represent mean values (n=3)

**Figure 5-figure supplement 1.**
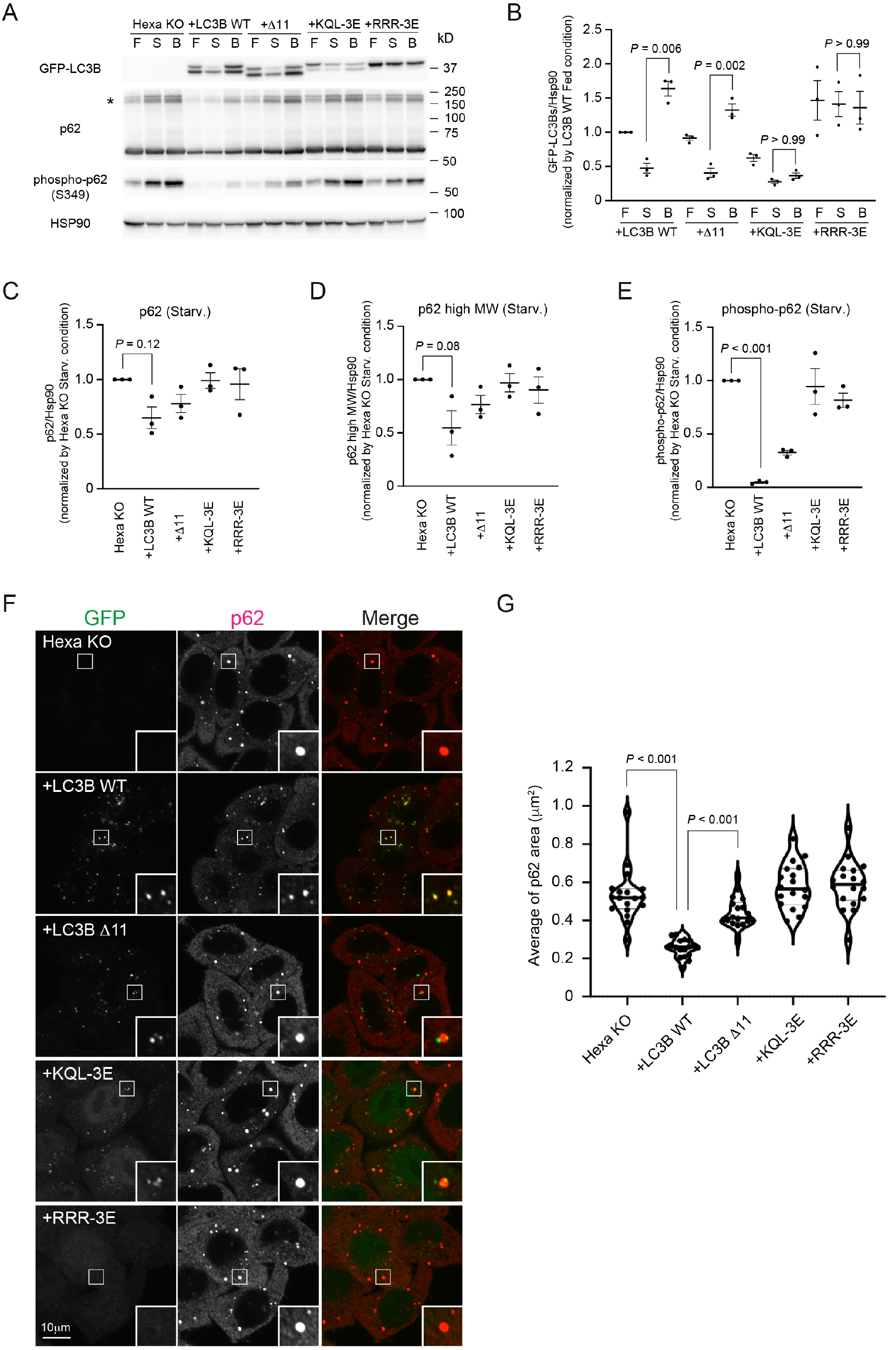
Membrane insertion of LC3B N-terminus contributes to efficient degradation of p62 body. **(A)** Hexa KO cells stably expressing GFP-LC3B WT, Δ11, KQL-3E or RRR-3E were starved for 8 h with (B) or without 100 nM Bafilomycin A_1_ (S), or cultured in full media (F). Cell lysates were analysed by immunoblotting using the indicated antibodies. The asterisk indicates the position of high molecular weight forms of p62. **(B-E)** Band intensity quantification of GFP-LC3Bs **(B)**, p62 **(C)**, p62 high molecular weight forms **(D)**, and phosphorylated p62 (Ser349) **(E)**. All data were normalized with those of HSP90. Data represent the mean ± SEM of three independent experiments. **(F)** The cells were starved for 2 h before p62 (red) was visualized. Scale bar, 10 μm. **(G)** The violin plot of average p62 area. The thick and thin lines in the violin plot represents the median and quartiles, respectively. n = 18. Differences were statistically analysed by one-way ANOVA and Turkey multiple comparison test.

**Table S1.**
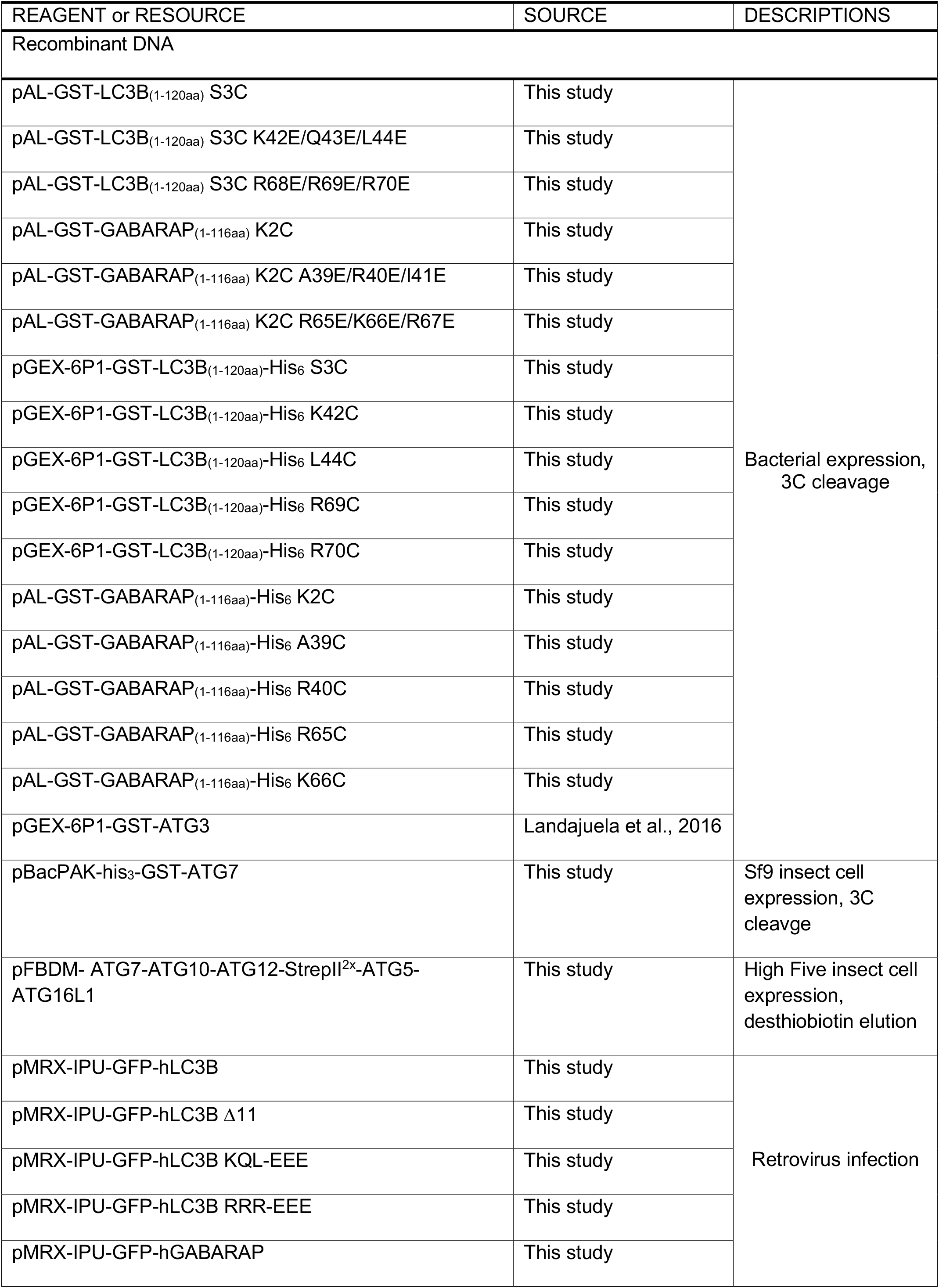
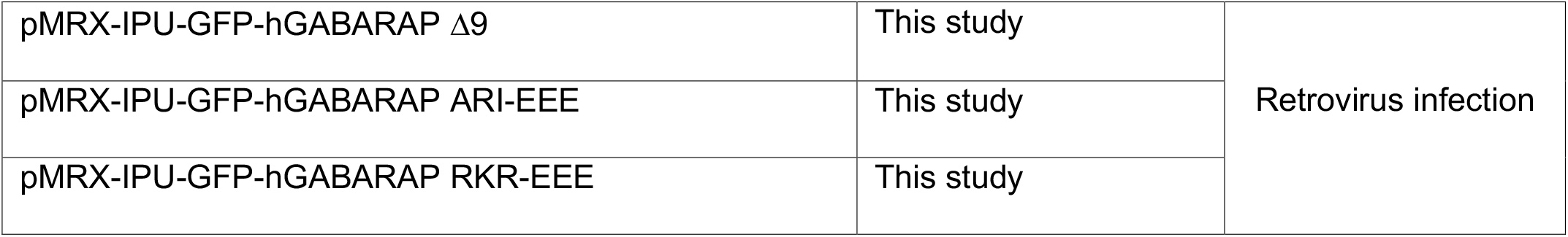
Constructs used in this study.

